# Early Spatiotemporal Deficits Become Competitive Advantages During Collective Expansion

**DOI:** 10.64898/2026.05.21.726850

**Authors:** Emma Dawson, Aishwarya Ganesh, Minsu Kim

## Abstract

Rapid spatial expansion into new territory is fundamental to population persistence, ecological dominance, and evolutionary success. Every race for space is initiated by founding populations, yet how their initial spatiotemporal conditions shape the outcome remains unclear. Here, by combining nonlinear theory with quantitative microbial experiments, we show that collective motility can overturn the classical logic of priority effects. Populations expanding by individual motility retain the expected advantage of an earlier start or a larger initial footprint. By contrast, under collective motility, nonlinear density-dependent feedback enables initially lagging populations to accelerate, transforming early deficits into long-term competitive advantage. Optimization predicts that this advantage is maximized when the delay scales log-linearly with initial density. We validate this scaling behavior across diverse swarming microbes, revealing a general regulatory strategy for competitive range expansion. Competition and evolution experiments further show that this behavior is adaptive and robustly preserved. Together, our results establish a quantitative framework linking the initial organization of founding populations to final spatial outcomes, helping explain how organisms regulate early dynamics to shape ecological success in space.

**Significance:** Rapid expansion into new territory often decides who will dominate. To understand this contest, we focused on how the race begins. We built a theoretical framework linking the initial conditions of founding populations to long-run competitive outcomes, then tested its predictions experimentally in microbial populations by controlling inoculum, quantifying range expansion, competing strains head-to-head, and performing laboratory evolution. Our results show that history is not merely the starting point of range expansion, but part of the regulatory strategy for winning space. This work provides a mechanistic framework for quantitatively thinking about range expansion across systems, from microbial colonization and biological invasions to infection spread, cancer metastasis and developmental migration.

## Introduction

When populations grow within a confined environment, they eventually exhaust local resources, limiting further growth. Range expansion provides a critical escape from these limitations across biological scales from microbes to insects, plants, and animals [1–3]. By spreading into new territory, organisms can avoid crowding, access untapped resources, and sustain population growth. Species that reach new habitats faster prevail, while slower expanders risk exclusion or even extinction [4]. This competition for space has far-reaching consequences across ecosystems, including biological invasion, biodiversity [5–8], and the spread of diseases [9]. Yet, despite its broad importance, the regulatory mechanisms organisms employ to win this race for space remain poorly understood.

Classical ecology predicts that early advantage is preserved [10]. Populations with larger initial footprints and expanding sooner maintain their positional lead over time. This maintenance of early advantage is known as priority effects, which are supported by diverse empirical observations. For example, early establishment can facilitate plant invasions by pre-empting space [11]. In animal populations, early arrivals to newly available habitat can benefit from elevated prey availability [12]. In human evolutionary history, populations at expansion wave fronts have been shown to contribute disproportionately to contemporary gene pools [13, 14].

In many biological systems, motility requires or is enhanced by interactions among neighboring individuals, a phenomenon known as collective motility [15, 16]. Such coordination can arise via chemical signaling, mechanical coupling, or hydrodynamic interactions [17, 18]. Collective motility is widespread across biological scales: bacteria swarm across solid surfaces [18–27], epithelial cells migrate collectively during wound healing and cancer metastasis [28–31], and animals such as birds, fish, and mammals exhibit coordinated group motion [16, 32–34].

In collective motility, population expansion cannot be approximated as the sum of independent individuals’ trajectories. Instead, individuals interact with their neighbors, and these interactions couple their movements. As density increases, motility can be enhanced, leading to faster redistribution of the population. This redistribution, in turn, reshapes the local density landscape, feeding back on the movement itself. Therefore, collective motility fundamentally alters the dynamical rules governing population expansion by introducing nonlinear density–motion feedback.

Nonlinear dynamics are often sensitive to initial conditions: small differences in the starting configuration can be amplified over time, producing divergent outcomes: a phenomenon sometimes referred to as the butterfly effect [35]. Feedback can either amplify or suppress this divergence [35, 36]. From an ecological perspective, this implies that biological history may exert a lasting influence on the dynamics of range expansion. If differences established early persist over long times, they effectively constitute a form of ecological memory. This also raises an intriguing possibility that populations do not merely inherit their initial conditions but may actively regulate them to maximize the competitive outcome during range expansion.

To understand how biological history affects this competitive outcome, we developed a theoretical framework linking the initial spatiotemporal organization of founding populations to long-term expansion. We tested its predictions in microbial populations, whose range expansion can be measured with high precision and whose initial geometry, timing, and density can be tightly controlled experimentally. Moreover, competitive consequences can be evaluated directly through head-to-head competition assays and laboratory evolution.

Bacteria provide an especially useful system because they exhibit both collective and individual motility. Their collective surface motility known as swarming is driven by coordinated movement of multicellular groups [37]. *Proteus mirabilis* is a canonical model system for bacterial swarming [38, 39]. Its advancing front can be quantified, enabling direct measurement of expansion dynamics over time (Fig. 1A) [40]. By contrast, when immersed in liquid or soft agar, *P. mirabilis* cells move via swimming, where individual cells move independently. This duality allows the consequences of collective versus individual motility for range expansion to be studied within a single organism.

**Figure 1.**
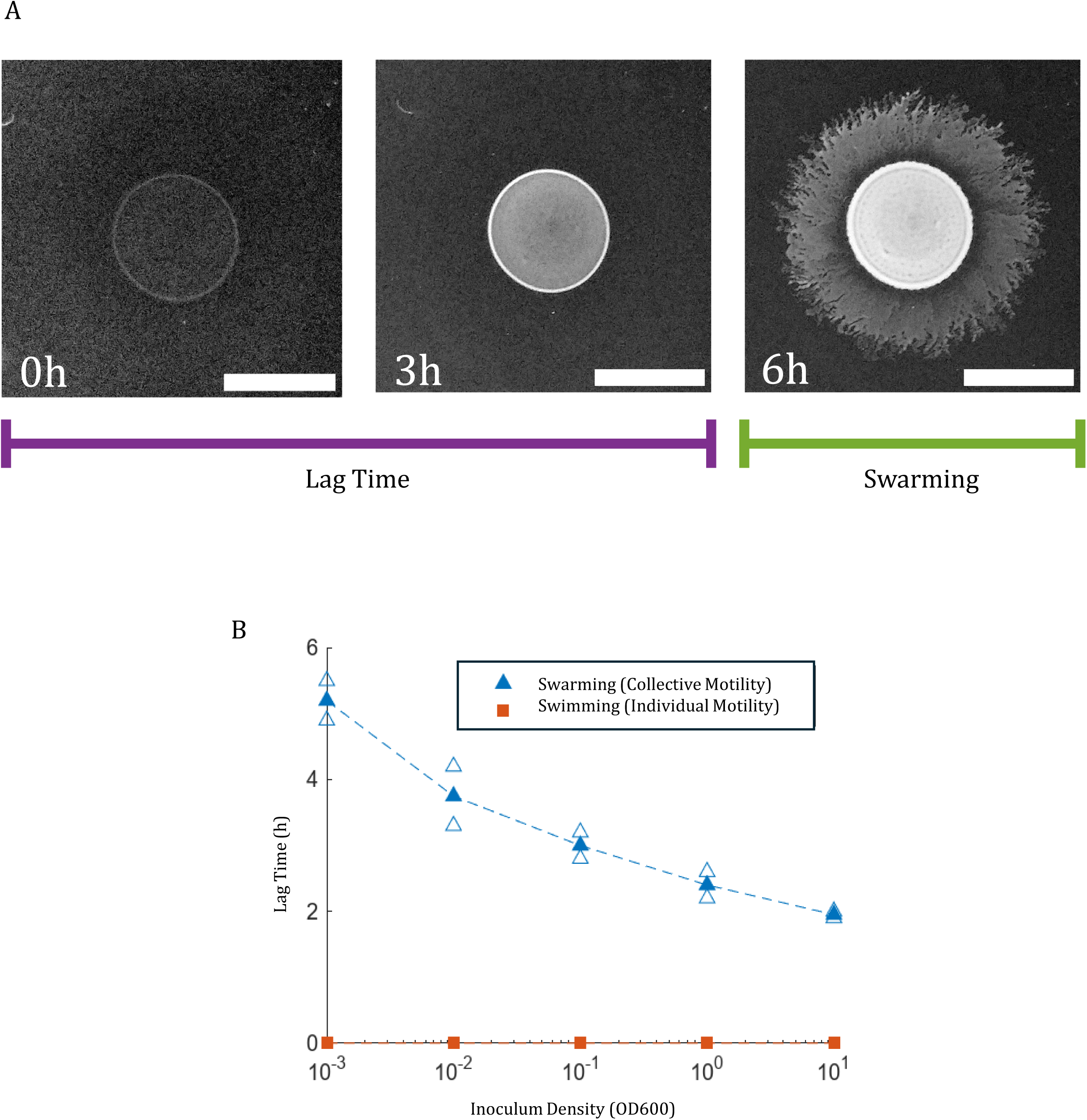
Density-dependent lag in collective expansion. (A) Representative timelapse images show the early stages of colony development for a *P. mirabilis* colony grown on an LB Agar plate. During lag time, the colony grows but does not expand. Once swarming begins, the colony expands its border radially. Scale bar 1cm. (B) Lag time as a function of inoculum density (OD_600_). Under swarming (collective motility; blue triangles), colonies exhibit a pronounced density-dependent delay prior to radial expansion. In contrast, swimming colonies (individual motility; orange squares) initiate expansion immediately, with no detectable lag across the densities tested. Two biological repeats were performed. Open symbols denote individual replicates; filled symbols denote the mean.

We found that *P. mirabilis* regulates its physiology to delay the onset of swarming expansion, whereas no such delay occurs during swimming. Our theoretical model reveals a nonlinear feedback mechanism by which collective motility transforms an initial spatiotemporal disadvantage into a net benefit during range expansion. Optimizing this mechanism predicts a log-linear relationship of delay with initial density, which we experimentally confirm across swarming bacteria. Competition assays between strains that either exhibit or lack this regulation, together with laboratory evolution for faster expansion, further show that this behavior is adaptive and evolutionarily robust. Together, these results reveal how the early history shapes range expansion, identifying regulatory mechanisms that populations employ to promote competitive success during spatial colonization.

## Results

### Density-dependent delay in bacterial swarming

We observed *P. mirabilis* swarming on an agar plate and tracked the colony edge (Fig. 1A). When we deposited cells at the center (Fig. 1A, leftmost image), the inoculum exhibited a lag phase in expansion, which was subsequently followed by rapid radial expansion (Fig. 1A and Supplementary Fig. 1). Interestingly, we found that the duration of this delay depends on inoculum density (Fig. 1B, blue color): colonies starting at lower initial density exhibited longer lag times before initiating expansion, yielding a monotonic inverse relationship on a log-linear plot.

For comparison, we examined its expansion in liquid, where cells move individually by swimming. Under these conditions, colonies initiated radial expansion immediately after deposition, with no detectable lag phase across the range of inoculum densities tested (Fig. 1B, orange color). Therefore, the density-dependent lag is a specific feature of swarming, but not swimming.

Bacterial swarming often involves differentiation into elongated, hyperflagellated cells [18, 21]. During this differentiation period, which typically lasts a couple of hours, colonies exhibit little or no outward movement [41–45], which explains the short lag period we observed at high inoculum density (Fig. 1B). However, this period of differentiation cannot explain the markedly prolonged lag observed at lower inoculum densities (Fig. 1B).

Consistent with this, our additional measurements indicate that the prolonged lag reflects active regulation. FlhDC is a master protein for swarming [46–48]. Although the factors regulating its expression are not fully resolved, its upregulation is required for swarming initiation while its downregulation inhibits swarming. At high inoculum density, *flhDC* expression increased soon after deposition (Supplementary Fig. 2). In contrast, at lower initial density, *flhDC* expression remained low for extended periods of time and then rose sharply at a later time point. In both cases, the increase in *flhDC* expression was immediately followed by swarming expansion (Supplementary Fig. 2). Together, the density-dependent timing of *flhDC* upregulation indicates that swarming initiation is controlled by an actively regulated decision point.

### Delayed initiation is costly under density-independent motility

At first glance, a prolonged lag period prior to expansion appears counterproductive. To test this intuition, we analyzed the Fisher–Kolmogorov equation, a classical reaction–diffusion model for population spread; see Supplementary Text, Section 1 for details. For compact initial conditions, as employed in our experiments, prior work has shown that solutions converge to a travelling-wave regime in which a fixed front profile propagates outward at a constant velocity [49, 50].

Here, we focus on how delayed initiation affects the expansion dynamics. To explicitly incorporate delayed initiation, we set the diffusivity to zero for a prescribed time window. During this lag phase, the biomass increases locally but did not spread spatially. This lag phase appears as an initial plateau in the colony radius–time plot (Fig. 2A). After the lag, motility was restored and expansion proceeded normally. In all simulations, colonies were initialized with identical total biomass for fair comparison; only the lag duration was varied.

**Figure 2.**
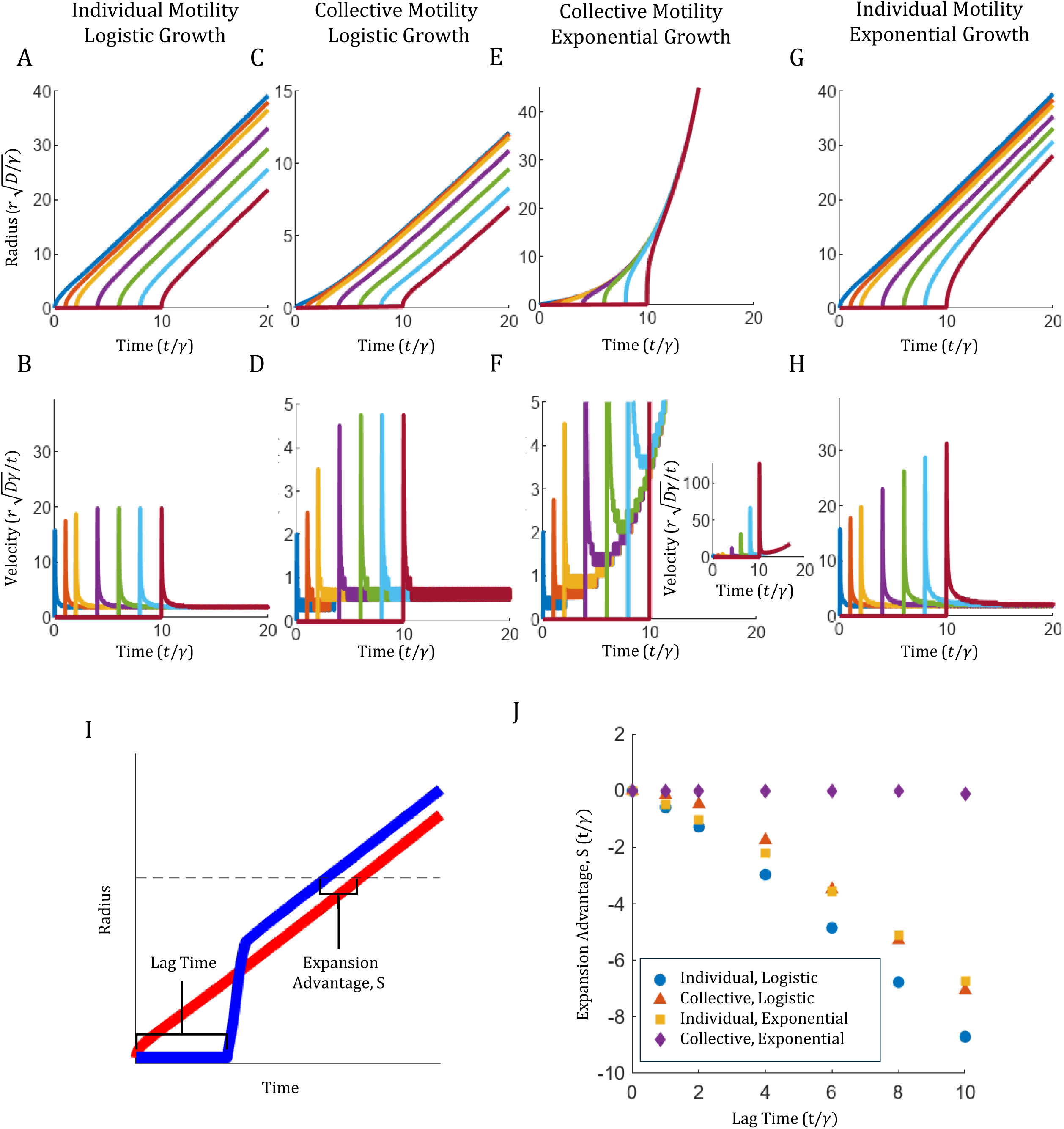
Simulations reveal how collective motility and growth regime determine the cost of delayed initiation. (A,B) Individual motility with logistic growth. See Supplementary Text Section 1 for model details. All quantities are plotted in dimensionless units as indicated in axes, following the definitions in the Supplementary Text. The lag phase appears as an initial plateau in the colony-radius trajectories (A) and as a corresponding delay in the front-velocity profiles (B). Colonies subjected to increasing lag times (colored curves) converge to travelling-wave solutions with identical slopes but retain fixed spatial offsets; delayed colonies remain permanently behind. See Supplementary Video 1. (C,D) Collective motility with logistic growth. Density-dependent transport partially mitigates the cost of delay but does not eliminate it; colonies still exhibit persistent radius offsets. See Supplementary Text Section 2. See Supplementary Video 2. (E,F) Collective motility with exponential growth. Delayed colonies exhibit faster initial velocities, with the velocities increasing with delay. Inset, full velocity trajectories over the entire range are shown. All trajectories ultimately converge, eliminating long-term spatial offsets. See Supplementary Text Section 4 and Supplementary Video 3A. Supplementary Video 3B shows a colony with longer lag phase. (G,H) Individual motility with exponential growth. Delayed colonies retain a persistent spatial disadvantage despite exponential growth. See Supplementary Text Section 3 and Supplementary Video 4. (I) Definition of expansion advantage, *S*. *S* is defined as the difference in arrival time at a fixed radius between a no-delay baseline colony (red) and a colony with lag (blue). Positive *S* indicates that delayed initiation yields a net benefit; negative *S* indicates a cost. (J) Expansion advantage as a function of lag time. Under logistic growth, delay remains costly for both individual and collective motility. Under exponential growth, delay becomes neutral with collective motility, whereas it remains penalized with individual motility.

Simulation results revealed that all colonies ultimately converged to travelling waves, expanding with a constant velocity (Supplementary Video 1). However, colonies with longer lags consistently remained spatially behind those that initiated expansion earlier (Fig. 2A). The corresponding instantaneous front-velocity profiles were similar apart from a temporal shift imposed by the lag (Fig. 2B), explaining why delayed colonies do not catch up. Once the travelling-wave regime was reached, this disadvantage was permanently locked in (Fig. 2A). Thus, within the classical Fisher–Kolmogorov framework, any delay prior to expansion is intrinsically costly and cannot be recovered at long times.

### Collective motility reduces the cost of delayed start

This raises a natural question: why would populations delay the onset of expansion? The classical Fisher–Kolmogorov framework assumes constant diffusivity, implicitly treating motility as an independent property of individual cells. Our above results demonstrate that under this assumption, delaying the onset of dispersal is always costly.

A lag phase, however, is not merely a period of inactivity, but reshapes the initial conditions for dispersal. By remaining stationary while growing, a colony accumulates biomass within a compact region and thereby increases local density. More importantly, swarming is a collective mode of motility in which effective transport increases with local cell density. The elevated density generated during lag phase may accelerate expansion. We therefore asked whether, under collective motility, investing in a lag phase could yield a net expansion benefit.

To test this hypothesis, we incorporated collective motility into the Fisher–Kolmogorov equation by allowing the diffusivity to increase with local population density, *D*(*u*), (Eq. S4; Supplementary Text Section 2). To model a lag phase, diffusivity was initially set to zero for a prescribed period and restored thereafter. For numerical calculations, we adopted the minimal form of density dependence in which diffusivity depends linearly on density. However, as will be evident in the theoretical analysis below, the conclusions do not rely on this specific linear form, but only on the general requirement that diffusivity increase with density, that is, *D*^′^(*u*) >0.

Our numerical calculations reveal that all colonies ultimately converged to travelling-wave solutions with identical propagation velocities (Supplementary Video 2 and Fig. 2C). Instantaneous front-velocity profiles reveal that colonies with longer lags exhibit transient velocity peaks slightly higher immediately after the onset of expansion (Fig. 2D). However, this enhancement is transient and weak: the density profiles soon relax toward the common travelling-wave form and the front velocities drop to the same low steady level (Fig. 2D). Colonies subjected to longer lag phases still entered the travelling-wave regime with smaller radii (Fig. 2C). Therefore, delay remains costly.

### Delay becomes neutral in the low-density regime

To understand why the cost of delay does not vanish under collective motility, we examined population growth dynamics. In logistic growth, the per-capita growth rate decreases linearly with cell density (Eq. S4). Spatially compact population profiles are therefore intrinsically penalized: for a fixed total biomass, confining the population to a smaller radius increases local density and thus reduces net growth (Supplementary Text, Section 2, Eq. S11). As a result, delayed colonies that remain spatially compact during the lag phase accumulate total biomass more slowly (Supplementary Fig. 3), leading to slower expansion (Supplementary Text, Section 2, discussion below Eq. S11). This density-dependent growth penalty is inherent to the functional form of logistic growth.

By contrast, early colonization of empty space typically begins from a small founder population. Under these low-density conditions, this growth penalty is negligible and population growth is therefore well approximated as exponential. To directly demonstrate that this strict penalty imposed by logistic growth is responsible for the persistent cost of delayed initiation, we replaced the logistic growth term with an exponential growth term in our model. Under this condition, simulations reveal a different outcome. Although colonies with lag phases begin expansion later, they exhibit very large transient velocity peaks immediately after the onset, which allow them to fully catch up to earlier-starting colonies (Fig. 2E,F and Supplementary Video 3A and 3B). Comparison of Supplementary Video 3A and 3B shows that longer lag phases produce stronger transient front acceleration, enabling complete catch-up regardless of lag duration. Importantly, this catch-up still requires density-dependent motility: even in this exponential growth regime, if motility is density-independent, delayed colonies fail to catch up (Fig. 2G-H and Supplementary Video 4). Thus, in the low-density regime, collective motility eliminates the long-term spatial cost of delay.

To summarize these findings, we next quantified the effects of delay under different growth and motility regimes. Colonies with different delays ultimately converge to the same velocity, differing only by a constant radius offset. Consequently, any advantage or disadvantage of delayed initiation can be evaluated by relative arrival times at a fixed spatial scale. Using a colony with no imposed waiting period as a baseline, we defined an expansion advantage, *S*, as the difference in arrival time at a target radius between the baseline colony and a colony that exhibited a delay (Fig. 2I). Positive values of *S* would indicate that the delayed colony reached the target radius earlier (a net expansion advantage), whereas negative values would indicate later arrival (a net cost of delay).

Using this definition, we analyzed how *S* depends on lag time across the different motility and growth regimes (Fig. 2J). Under individual motility, *S* becomes increasingly negative with longer lag, showing that delay produces a persistent spatial disadvantage, recapitulating the classical priority-effects. By contrast, under collective motility in the low-density regime, *S* remains zero (purple diamond), again showing that delay carries no long-term spatial cost.

### Reformulating temporal delay into a geometry problem theoretically confirms numerical predictions

This result can also be tested through an equivalent formulation based on initial geometric organization. In the delay scenario, growth proceeds locally while spatial spreading is suppressed, allowing biomass to accumulate without expansion. This process produces a spatially compact population at the onset of motility. An equivalent state can be generated without an explicit delay by initializing colonies with the same total biomass but different initial radii.

This reformulation utilizes the fact that, once expansion begins, the governing equations depend only on the spatial distribution of biomass (and the total biomass), not on how that distribution was generated. A colony initialized with a smaller radius represents the effective end state of a population that has grown locally without spreading, whereas a colony initialized with a larger radius corresponds to a population whose biomass has already been distributed over space by earlier spreading.

This analogy can be also explained by the radius–time trajectories. In the delayed-initiation protocol, a colony that exhibits a delay remains stationary initially and then begins expanding. As a result, at any fixed observation time after expansion has begun (but prior to the convergence), colonies with different delays occupy different radii while having the same total biomass. In other words, delayed initiation naturally generates a family of states that differ only in their spatial extent at a later common time. Thus, analyzing colonies with equal biomass but different initial radii is mathematically equivalent to taking a snapshot of the delayed-initiation dynamics at a later time and using that snapshot as a new initial condition. Consistent with this equivalence, our numerical simulations show that colonies with different initial radii fully catch up under collective motility, converging to the same long-time expansion trajectory (Supplementary Fig. 4), as we have shown for colonies with the same radii but delays (Fig. 2E).

An advantage of this geometric reformulation is that it renders the problem analytically tractable (Supplementary Text, Section 5), providing an intuitive understanding. The analytical solutions reveal that under individual motility, two colonies initialized with identical biomass but different radii approach a fixed spatial separation at long times, with the separation determined by the initial radius (Eq. S39). Thus, the early disadvantage is preserved in the long-time limit.

By contrast, under collective motility, there exists a unique asymptotic travelling-wave solution (Eq. S47). This solution contains a single parameter equal to the total initial biomass, and does not depend on the initial radius, indicating that the asymptotic state does not retain geometric memory of how the biomass was initially distributed. Therefore, both numerical simulations and theoretical analyses predict that under collective motility, long-time expansion becomes insensitive to the initial spatio-temporal organization of founding populations.

Our theoretical derivation also provides mechanistic insight into this insensitivity. Collective motility implies that effective diffusivity is an increasing function of local density, *D*(*u*), with *D*^′^(*u*) > 0. For a fixed total biomass, distributing cells over a smaller radius produces a more compact profile with higher density. This is expected to accelerate the expansion, as we have seen in numerical simulation (Fig. 2F).

This argument can be made more explicit by the model structure. Rewriting the Fisher–Kolmogorov equation with density-dependent diffusivity introduces an additional transport term that is mathematically equivalent to an advective contribution in a convection–diffusion equation (Supplementary Text, Section 2). This term generates a self-induced transport with velocity proportional to *D*^′^(*u*) ∂*u*/ ∂*x* (Eq. S5-6) which augments the diffusive flux. For collective motility, *D*^′^(*u*) >0, thus this contribution is outward. Moreover, for a fixed total biomass, a more compact colony exhibits steeper density gradients ∂*u*/ ∂*x*. These two factors multiply to create a strong outward advective flux, accelerating expansion upon the onset of expansion. This mechanism does not depend on the specific functional form of *D*(*u*), only on the condition *D*^′^(*u*) > 0, that is, collective motility. The resulting outward bias persists until the spatial profile relaxes to match the no-delay reference colony, enabling complete catch-up.

### Experiments validate the predicted convergence of spatial trajectories

An additional advantage of reformulating the originally temporal problem as one of initial geometry is that inoculum geometry can be tuned experimentally, enabling a direct test of the theoretical prediction. To isolate the effect of initial geometry from the density-dependent lag program inherent to the wild-type strain, we utilized a previously isolated mutant often referred to as precocious or simply “Prec” in the literature [47]. In this strain, deletion of a regulatory region in the *flhDC* promoter relieves repression, resulting in immediate upregulation of *flhDC* expression upon inoculation [47].

Our experiments further revealed the absence of density-dependent lag in this mutant. We inoculated two Prec colonies with the same initial biomass but different initial radii. Representative time-lapse images show that while these colonies exhibited a brief lag corresponding to the differentiation period, this lag was identical across inoculum radii, leading to a common onset of expansion (Fig. 3A). This feature allows us to examine the effect of initial geometry without the confounding influence of density-dependent variation in expansion onset in the wild-type strain, enabling a direct test of our theory.

**Figure 3.**
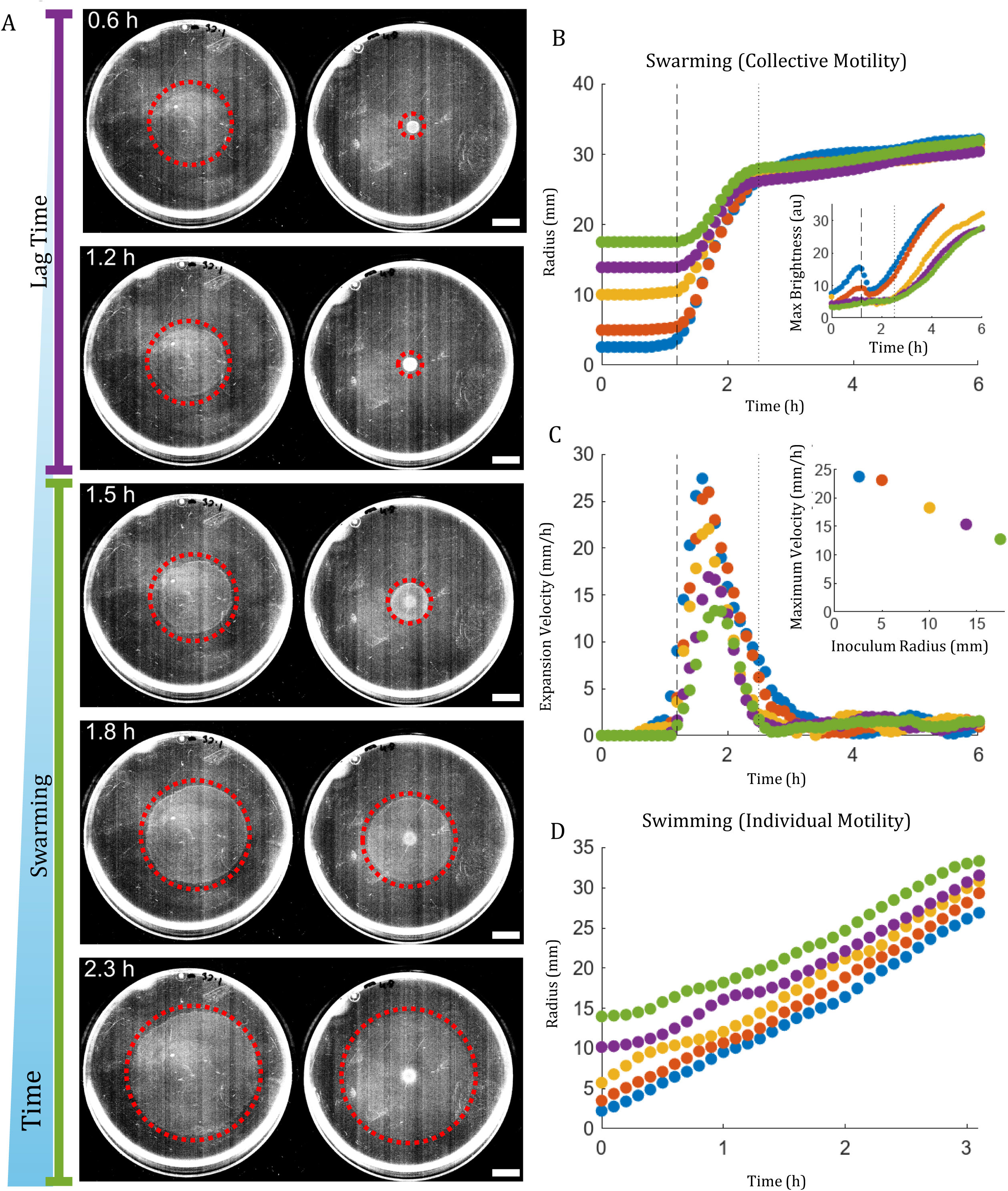
Experiments validate the predicted convergence of spatial trajectories under collective motility. (A) Representative time-lapse images for the two different initial radii but the same initial biomass are shown in Fig. 3A. The dotted red circle is guide to mark the colony front. Once swarming begins, the smaller colony expands more quickly than the larger colony and quicky catches up to the larger colony. Scale bar is 1 cm. (B) Swarming (collective motility). Colonies initiated from more compact geometries exhibit pronounced catch-up dynamics, rapidly reducing initial radius differences and converging toward similar long-time expansion trajectories. Vertical dashed lines indicate the onset of expansion. Dotted line is when they catch up. Biological repeat is shown in Supplementary Fig. 5A. Inset: Maximum brightness (proxy for biomass) as a function of time. Initially, the brightness increases while the colony grows during lag phase. At the onset of expansion (dashed line), the biomass is spread over a larger area, causing a dip in maximum brightness. After the initial dip, max brightness continues to grow past the time of radii convergence (dotted line), indicating that colonies remain well below carrying capacity during the catch-up period. (C) Expansion velocity for swarming colonies. The expansion velocity early after swarming (during the catch-up period) is higher for colonies that start out more compact, and at higher density. Inset: Maximum expansion velocity observed for each colony shows a clear dependence on inoculum radius. (D) Swimming (individual motility). Colonies were initiated with identical total biomass but different initial radii. During expansion, colonies maintain their initial spatial offsets: smaller-radius inocula remain persistently behind larger-radius inocula, indicating that individual motility preserves geometric memory. Biological repeat is shown in Supplementary Fig. 5B.

We found that, after the common onset of swarming, the colony initialized with a smaller radius (Fig. 3A, right) expanded markedly faster than that initialized with a larger radius (left), rapidly closing the initial radius gap. The two colonies ultimately converged toward the same radius at later times.

We further tested this convergence across a broader range of initial radii by tracking the colony radius over time (Fig. 3B and Supplementary Fig. 5A). Despite their different initial radii, all colonies started expanding at the same onset time (dashed line), consistent with our observation above. In every case, colonies converged toward the same long-time expansion trajectory (dotted line, Fig. 3B), experimentally validating our prediction that long-time expansion is indeed insensitive to initial geometry.

Next, we determined the intensity signal within colonies as a proxy for population growth (Fig. 3B, inset). At the time of the convergence (dotted line), colonies were still actively growing, with no evidence of density-limited saturation. This justifies our model assumption of exponential growth during the catch-up phase.

Our theoretical analysis attributed this geometric insensitivity to a stronger transient velocity boost in spatially compact colonies immediately after swarming onset. To test this prediction experimentally, we quantified the front velocity (Fig. 3C). Colonies initialized with smaller radii indeed exhibited larger transient peaks in expansion velocity (Fig. 3C). Consistent with this, the maximum front velocity, corresponding to the peak value (Fig. 3C inset), was higher for colonies with smaller initial radii, supporting the theoretical prediction.

A second key prediction of the theory was that this catch-up mechanism depends specifically on collective motility. To test this, we repeated the same experiment under swimming conditions, where motility is individual and no longer density-enhanced (Fig. 3D; Supplementary Fig. 5B). In this case, colonies with different initial radii expanded approximately in parallel and retained their initial spatial offsets throughout the experiment, with no evidence of catch-up.

Together, our numerical, theoretical, and experimental results show that collective motility suppresses the initial geometric imprint of the founding population by generating a strong transient expansion boost in spatially compact colonies. As a result, long-time expansion becomes largely insensitive to initial spatiotemporal organization. By contrast, under individual motility, the initial imprint is preserved and continues to shape the expansion trajectory.

### A biological constraint unlocks a benefit to delayed initiation

Our results with the nonlinear expansion theory show that delayed initiation can carry no inherent long-term cost under collective motility. However, this theoretical neutrality alone does not explain why swarming bacteria biologically regulate the onset of expansion in the first place. To address this, we next examined the basic physiology of resource allocation. In our simulations, we assumed an identical growth rate, *λ*, regardless of whether cells were motile. In reality, motility is energetically and biosynthetically costly: diverting resources to the synthesis and operation of flagella and motors is well known to reduce cellular growth rates [51–53]. Such growth–motility trade-offs have been widely documented across a wide range of organisms, indicating that they are a fundamental property [54, 55]. Consistent with this expectation, our experiments confirm a similar trade-off in *P. mirabilis*; a motile strain (the precocious strain above) exhibits a lower growth rate than a non-motile strain (Δ*motB*) under otherwise identical conditions (Fig. 4A).

**Figure 4.**
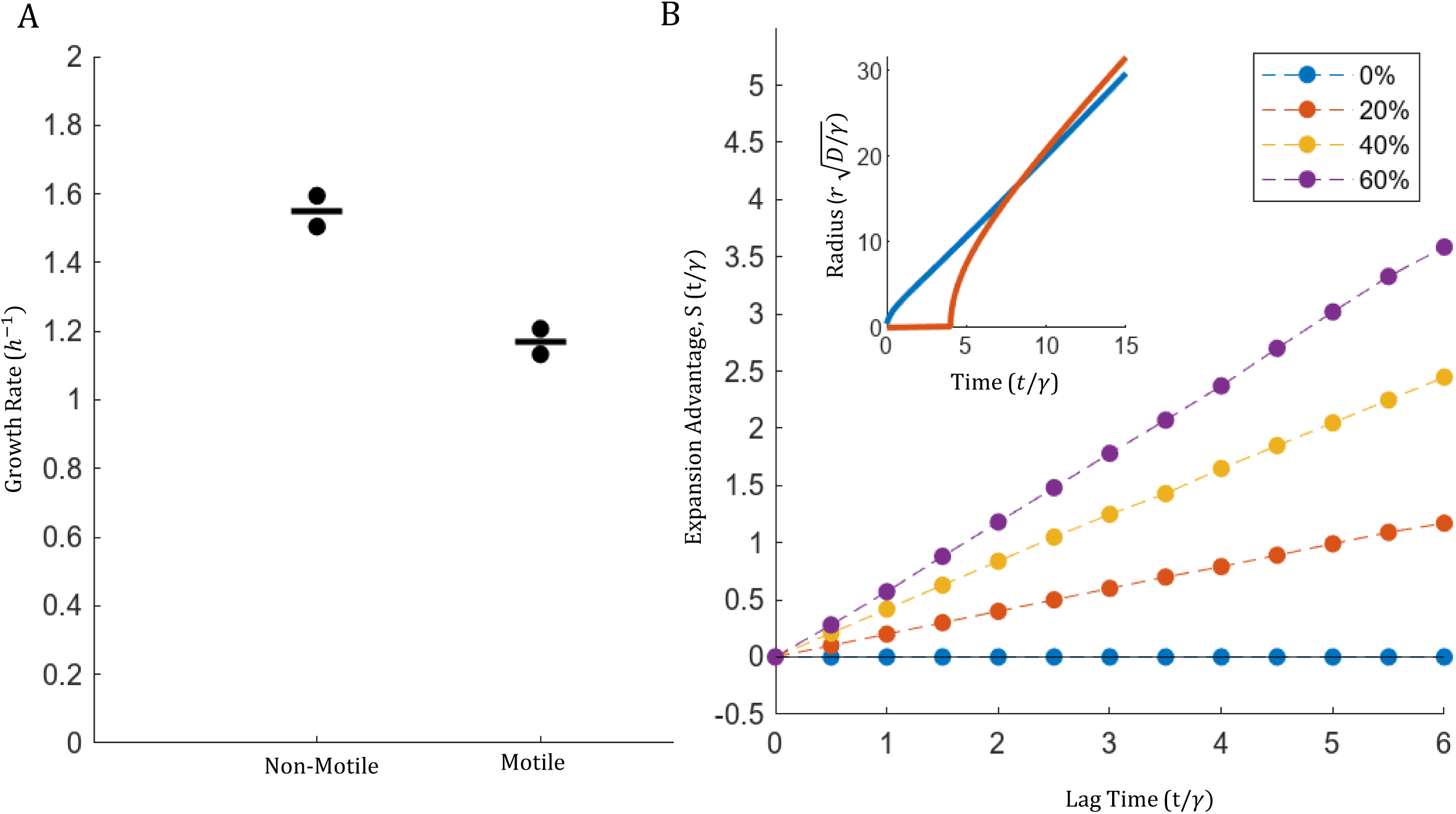
A growth–motility trade-off converts delay into competitive advantage. (A) Growth rate measurements for non-motile (Δ*motB*) and motile (Prec) strains. Motile cells exhibit slower growth rates, demonstrating a growth–motility trade-off. Points represent biological replicates; horizontal bars indicate the mean. (B) Expansion advantage *S* as a function of lag time under increasing growth-rate benefit during the non-motile phase (0%, 20%, 40%, 60%). When growth during the lag equals that during motility (0%), delay remains neutral. As the growth advantage during the lag increases, delayed colonies accumulate additional biomass and ultimately gain a net expansion advantage ( *S* > 0). Inset: Representative radius–time trajectories illustrating overtaking dynamics when growth during the lag exceeds that during motility.

Motivated by this growth–motility trade-off, we modified our model to allow cells to grow at a higher rate, *λ*_lag_ > *λ*, during lag phase, when motility is suppressed. After the onset of expansion, growth proceeds at the reduced rate *λ*. Under this modified model, colonies with a delayed start can accumulate higher biomass during the lag phase, which enables them to jump-start their expansion to ultimately surpass the immediate start colonies (Fig. 4B, inset). Quantifying the net expansion advantage (*S*, as defined in Fig. 2I) reveals that the advantage increases monotonically both with the time invested in the lag and with the magnitude of the growth-rate advantage during the lag phase (Fig. 4B). Even an infinitesimal growth advantage is sufficient to make delayed initiation beneficial under collective motility (Fig. 4B and Supplementary Fig. 6).

### Existence of a nontrivial minimum predicts expansion initiation at a critical density as an optimal strategy

Our results above show that under collective motility, delaying expansion can provide a competitive advantage. Yet, this analysis focused on the asymptotic travelling-wave regime, which effectively corresponds to expansion in unbounded space. In this limit, there is no intrinsic bound on the beneficial duration of lag: waiting longer continues to improve competitive performance. Natural habitats, however, are finite: populations need only spread a fixed distance to occupy the available area.

We therefore consider the total time to reach a finite boundary,

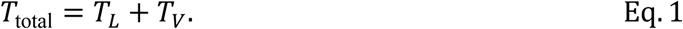

where *T*_*L*_ is the lag time and *T*_*V*_ is the subsequent travel time. On one hand, initiating expansion at a higher density requires a longer lag, which increases the total time to occupation and therefore incurs a direct temporal cost. On the other, a higher initiation density accelerates subsequent expansion and thereby reduces the travel time, providing a compensating benefit. In the race for space, the optimal strategy is therefore the one that minimizes *T*_total_.

We optimized this cost–benefit balance mathematically: see Supplementary Text, Section 6. Briefly, in Eq. 1, *T*_*L*_(*u*) is an increasing function of *u*, because reaching a higher initiation density requires more time. *T*_*V*_(*u*) decreases with *u*, because higher density enhances subsequent expansion. Therefore, the total occupation time *T*_total_(*u*) always has a global minimum (Supplementary Text, Section 6c). The trivial minimum is *u*^∗^ = *u*_0_. This corresponds to immediate initiation without delay, predicting the classical priority effects.

However, a nontrivial interior minimum, *u*^∗^ > *u*_0_, emerges if *T*_*V*_(*u*) decreases steeply with *u* while *T*_*L*_(*u*) increases only slowly with *u* (Supplementary Text, Section 6d). Biologically, these requirements correspond to collective motility, which makes expansion strongly density-enhanced, and a growth advantage during the lag phase, which reduces the temporal cost of waiting (Supplementary Text, Section 6e). Notably, these two are precisely the conditions considered in this study. The existence of a nontrivial interior optimum means that lagging until a critical density minimizes the total time required to occupy space, making threshold-like initiation an optimal strategy for the race for space.

### Experiments confirm critical initiation density as a universal feature of swarming

Motivated by this prediction, we asked whether *P. mirabilis* delays swarming until it reaches a critical density. Under this hypothesis, the lag time *T*_*L*_ should be determined simply by the time required for population growth to reach the swarming threshold. Assuming exponential growth during the lag phase,

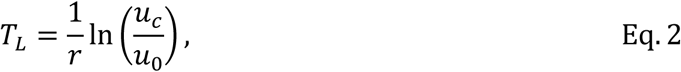

where *u*_0_ is the initial density, *u*_*c*_ is the critical density for swarming initiation, and *r* is the growth rate during lag. Because *u*_*c*_ is constant, this model makes a quantitative prediction that *T*_*L*_ should vary linearly with −ln *u*_0_ as the initial density is varied.

This prediction explains our earlier observation that *P. mirabilis* lag time decreases log-linearly with increasing initial density (Fig. 1B, replotted as Fig. 5A). Motivated by this consistency, we next measured the lag time of other swarming bacteria, including *Bacillus subtilis*, *Pseudomonas aeruginosa*, and *Escherichia coli*, all of which exhibited similar log-linear scaling (Fig. 5B–D).

**Figure 5.**
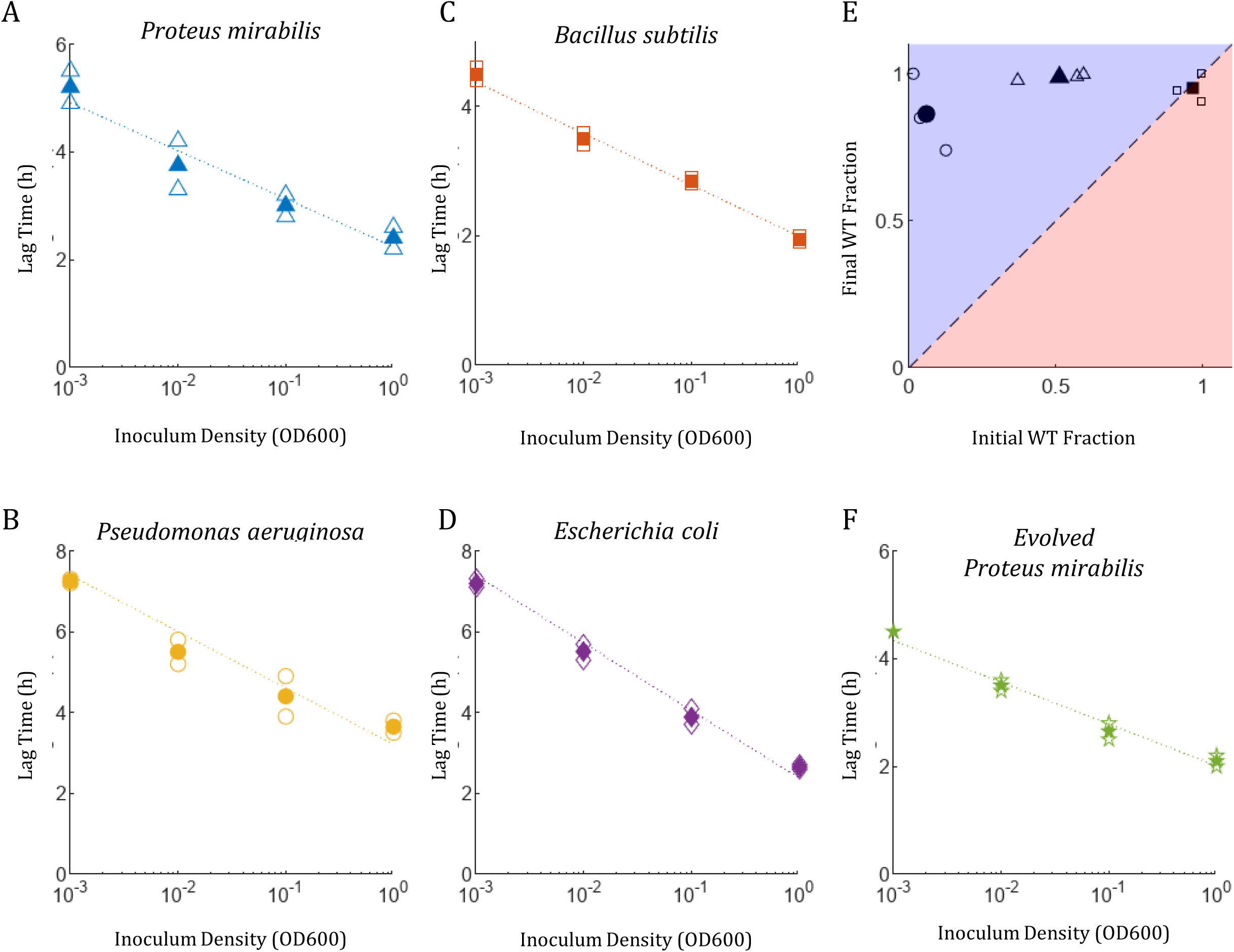
Threshold-like initiation is conserved and adaptive in the race for space. (A-D, F) Swarming lag time as a function of inoculum density for *Proteus mirabilis, Bacillus subtilis, Pseudomonas aeruginosa*, *Escherichia coli* and evolved *P. mirabilis*. Across all strains tested, lag time decreases monotonically with inoculum density and is well approximated by a log-linear relationship, which indicates that swarming is initiated at a critical density (Supplementary Fig. 7). Open symbols indicate individual replicates, closed symbols indicate the mean. (E) The WT:Prec ratio is represented as the fraction of WT cells in the total population. The x-axis indicates the initial WT fraction in the inoculum, and the y-axis indicates the final WT fraction after the mixed population swarmed across an LB agar plate. The diagonal dashed line marks neutrality, where the final fraction equals the initial fraction. Points above the diagonal (blue region) indicate that WT increased in frequency and therefore had a competitive advantage, whereas points below the diagonal (red region) indicate that Prec increased in frequency. Across a broad range of initial mixture ratios, we found that the final WT fraction remained above the neutral expectation, showing that WT outcompetes Prec during range expansion and can increase both when rare and when common. Open symbols indicate individual replicates, and closed symbols represent their mean.

The slope of this relationship is set by the lag-phase growth rate *r*, which we estimated by fitting each dataset (Eq. S61 and Supplementary Fig. 7A). Using these fitted values, we then directly inferred *u*_*c*_, i.e., the critical density that defines the onset of expansion (Supplementary Text, Section 6f). The inferred *u*_*c*_values were constant for each species regardless of the inoculum density (Supplementary Fig. 7). Together, these experimental results validate the predicted threshold-like initiation at a critical density as a general regulatory strategy of collective expansion.

### Threshold-like initiation is adaptive and evolutionarily robust in the race for space

In our model, threshold-like initiation emerges because it confers a competitive advantage in the race for space. To test this advantage experimentally, we conducted competition assays. We mixed the precocious mutant (Prec), which lacks the regulated lag, with WT at a 1:1 ratio, inoculated the mixture at the center of agar plate, and allowed it to swarm out. When the mixture arrived at the plate edge, we then sampled cells from the edge and determined the WT:Prec ratio. We found that WT became strongly enriched, reaching nearly 100% (Fig. 5E, triangle), indicating a clear competitive advantage. These results again support our prediction that regulating lag to reach a critical density is a beneficial strategy.

A trait is considered evolutionarily stable if it can increase when rare yet resist invasion when common [56]. To test this invasion-stability logic, we repeated the competition experiment while varying the initial WT:Prec ratio. When WT was initially common (99:1), Prec failed to increase, indicating that WT resists invasion (Fig. 5E, square). Conversely, when WT was initially rare (1:99), it became highly enriched after competition (Fig. 5E, circle), indicating that WT can invade a Prec population. Together, these results show that the regulated lag strategy of WT is stable against invasion by the early initiation strategy of Prec.

To further test the evolutionary robustness of the regulated lag, we subjected *P. mirabilis* to experimental evolution for faster range expansion via serial edge selection. A WT colony was inoculated at the plate center, allowed to swarm to the boundary, and cells from the advancing edge were transferred to the center of a fresh plate; this procedure was repeated every twelve hours. After 14 days of repeated transfer, the evolved population expanded roughly twice as fast as the WT ancestor (reaching the boundary of a 10-cm dish in 5.4 h compared with 11.5 h for WT). Despite this marked increase in swarming speed, the evolved strain retained the same log-linear scaling of lag time with initial density as the WT strain (Fig. 5F). Thus, selection for faster expansion did not eliminate density-dependent lag regulation, suggesting that this strategy is robust under continued selection for the race for space.

## Discussion

Rapid spatial expansion is critical for populations competing for space. In this work, we developed a theoretical framework linking the early spatiotemporal conditions of founding populations to their later competitive outcomes. For individual motility, a population that begins earlier keeps its lead, giving rise to classical priority effects [57, 58]. Yet in many biological systems, movement is not purely individual but instead emerges from coordinated interactions among neighboring individuals [15, 59]. We found that this collective motility fundamentally changes the role of biological history in range expansion. Nonlinear feedback between density and motility generates self-induced outward advection, producing a strong transient velocity boost that renders long-time expansion insensitive to initial spatial configuration. Numerical simulations show that colonies with identical total biomass converge to the same asymptotic solution regardless of their lag periods. By reformulating the temporal lag as a geometric problem, we derived analytical solutions that further demonstrate this insensitivity. In this regime, lag is no longer intrinsically costly. Consequently, any growth advantage conferred by growth–motility trade-offs turns lag into a benefit. Optimization predicts that this benefit is maximized when expansion begins after the population reaches a critical density.

We extensively tested this model in microbial populations by controlling the inoculum conditions, measuring the dynamics of range expansion quantitatively, competing strains against each other, and performing laboratory evolution. The observed log-linear scaling between lag time and initial density supports critical-density initiation across diverse swarming microbes. In *P. mirabilis*, wild-type cells outcompeted a precocious mutant that initiates expansion early, demonstrating the adaptive value of regulated lag. This scaling is robustly maintained under selection for fast expansion, even as evolved populations increased their swarming speed. Together, these experimental data strongly support our model, revealing a general adaptive strategy for competitive range expansion.

This finding provides an ecological perspective on molecular control of motility. Quorum sensing is a molecular regulatory mechanism in which signaling molecules accumulate locally and trigger coordinated behavior once a threshold concentration is reached [60]. Our results place this molecular control in the ecological context, providing a dynamical explanation for why quorum sensing so often regulates motility in bacteria [60, 61].

Collective motility corresponds to a regime in which motility is enhanced by local interactions, yielding an effective diffusivity that increases with density (*D*^′^(*u*) > 0). A useful opposite limit, though not analyzed here, is one in which mobility decreases with density (*D*^′^(*u*) < 0). Such behavior can arise from jamming, in which crowding and mechanical constraints hinder rearrangements and reduce mobility [22, 62]. Reversing the sign of *D*^′^(*u*) reverses the density-gradient-driven advective contribution, producing an inward bias. As a result, compact colonies would decelerate rather than accelerate. This will cause early differences to intensify, strengthening priority effects.

More broadly, our work elucidates how the initial conditions of founding populations shape long-term expansion dynamics. Initial conditions are not merely inherited; they can be managed. Organisms can regulate early spatiotemporal organization to win the race. These findings help us think about spatial competition across systems, including biological invasions, biofilm spreading, tumor metastasis, epithelial wound healing, and animal dispersal.

## Materials and Methods

### Strains and media

*P. mirabilis* ATCC 7022 and its derivatives were used (Supplementary Table 1). Liquid cultures were prepared in Luria-Bertani (LB) broth at 37°C with shaking at 200 rpm. An overnight culture was inoculated from -80°C frozen stock the day prior to each experiment and allowed to grow overnight. A small volume of the overnight culture was suspended in fresh LB broth and grown for at least two doublings before used in the experiments described in this text. Cell density was determined by quantifying the optical density at 600 nm wavelength (OD_600_) using a Genesys20 spectrophotometer (Thermo Fisher) with a standard cuvette (1.6100-Q-10/Z8.5; Starna Cells).

To produce inoculums of varied densities, an experimental culture was prepared in 60 ml of LB broth and grown to OD_600_=0.4. The culture was centrifuged, the supernatant was discarded, and the cell pellet was resuspended in 20 ul of pre-warmed, fresh LB. The culture was diluted to the desired densities in pre-warmed, fresh LB.

For swarming experiments, 16 ml of LB agar was solidified in a 10 cm Petri dish. The agar concentrations, chosen to facilitate swarming, were as follows: *Proteus mirabilis*, 1.25%, *Bacillus subtilis*, 0.8%, *Pseudomonas aeruginosa*, 0.5%, *Escherichia coli*, 0.4%. The LB agar plates were prepared on the same day as each experiment and allowed to rest at room temperature on the lab bench for 4 hours, before being moved to 37°C for pre-warming 30 min before inoculation.

For swimming experiments, 16 ml of LB with 0.5% agar was solidified in a 10 cm Petri dish. The LB agar plates were prepared on the same day as each experiment and allowed to rest at room temperature on the lab bench for 4 hours, before being moved to 37°C for pre-warming 30 min before inoculation.

### Imaging and image analysis

For swarming and swimming assays, colonies were imaged using a document scanner (Epson PerfectionV39 II) controlled by automation software (RoboTask Lite), following an imaging protocol developed by Bru et al. [63]. For swimming lag times, cells were imaged using an inverted microscope (Olympus IX83P2Z) and a Neo 5.5 scientific CMOS camera (Andor), controlled by MetaMorph software (Molecular Devices), and housed in an incubator (InVivo Scientific) at a temperature of 37°C. Images were analyzed using Manual Tracking, a freely available plugin for ImageJ, and custom MATLAB software.

### Swarming lag time experiments

Liquid cultures with OD_600_ 10, 1, 0.1, 0.01, and 0.001 were prepared as described above. For each density, a 4 µL droplet of culture was inoculated onto the center of a pre-warmed LB swarming agar plate and allowed to dry. Once dry, the plates were moved to an incubator at 37°C with a water reservoir to maintain humidity. Plates were imaged using a document scanner (Epson PerfectionV39 II), as described above.

### Swimming lag time experiments

Liquid cultures with OD_600_ 10, 1, 0.1, 0.01, and 0.001 were prepared as described above. For each density, slides were prepared as follows: a 22 mm × 22 mm glass coverslip was secured to a 3 in × 1 in glass slide (VistaVision) using double-sided tape (3M Scotch) along three of its edges, creating a small gap between the coverslip and slide the height of the double-sided tape. From the open edge of the coverslip, the gap was flooded with fresh LB, and the slide was prewarmed to 37°C. Just before use, each slide was mounted onto the microscope, and the excess LB was removed using a pipette. A 0.5 µL aliquot was deposited at the open edge of the coverslip, and cells were imaged as they swam through the gap.

### Swarming assay

The same number of cells was inoculated onto each swarming LB agar plate and spread to circles of varying radii using an inoculum loop. To facilitate even spreading, several cultures at different densities were prepared and inoculated in different volumes to keep the total number of cells constant. For each radius, the inoculum density and volume used are: 1 mm radius, 2 µL at OD_600_=16, 2 mm radius, 8 µL at OD_600_=4, 5 mm radius, 32µL at OD_600_=1 and for 10 mm and 15 mm radii, 64 µL at OD_600_=0.5. Once dry, the plates were then moved to an incubator at 37°C with a water reservoir to maintain humidity. The plates were imaged using a document scanner, as described above.

### Swimming assay

The same number of cells was inoculated onto each swimming LB agar plate and spread to circles of varying radii using an inoculum loop. To facilitate even spreading, several cultures at different densities were prepared and inoculated in different volumes to keep the total number of cells constant. For each radius, the inoculum density and volume used are: 1 mm radius, 2 µL at OD_600_=16, 2 mm radius, 8 µL at OD_600_=4, 5 mm radius, 32 µL at OD_600_=1 and for 10 mm and 15 mm radii, 64 µL at OD_600_=0.5. Once each plate was inoculated, an inoculum needle (for inoculum radius 1, 2, 5 mm) or inoculum loop (for inoculum radius 10, 15 mm) was used to gently mix the inoculated cells into the top of the swimming agar plate to a depth of 1-2 mm. The plates were then moved to an incubator at 37°C with a water reservoir to maintain humidity. The plates were imaged every 6 minutes using a document scanner, as described above.

### Swarming evolution experiments

To evolve the strain of *P. mirabilis*, we started with our Wild Type *P. mirabilis* ATCC 7022. At time zero, 4 µL of liquid culture with OD600 of 1 was inoculated onto the center of a pre-warmed swarming LB agar plate and allowed to dry. Once dry, the plate was moved to an incubator at 37°C with a water reservoir to maintain humidity. After 12 hours, the plate was removed from the incubator. Using a sterile spatula, a 5 mm ×15 mm section was excised from the edge of the colony, including the agar beneath it. This section of agar was added to a 1.5 mL microcentrifuge tube, and 100 µL of fresh, pre-warmed LB broth was used to gently wash the cells off the agar pad by pipetting. The density of the resulting cell culture was adjusted to an OD600 of 1 and 4 µL of the adjusted cell culture was inoculated onto a fresh, pre-warmed swarming LB agar plate and allowed to dry. Once dry, the plate was moved to an incubator at 37°C with a water reservoir to maintain humidity. This selection process was repeated every 12 hours for 14 days to generate the evolved strain used in this study.

### Surface growth rate experiments

For the surface growth rate experiments in Fig. 4A, swarming LB agar plates were prepared as described above. An exponentially growing liquid culture with OD_600_=0.1 was prepared, and a 100 µL aliquot was added to each plate and spread using 6 sterile glass beads per plate until dry. The glass beads were then removed, and plates were moved to an incubator at 37°C with a water reservoir to maintain humidity. Every hour, two plates for each strain were removed from the incubator for measurement. The plates were flooded with 5 mL of pre-warmed LB, and cells were gently dislodged from the surface using an inoculum loop. The LB and cells were collected from each plate and the OD_600_ of the suspension was measured.

### Swarming microscopy experiments

For swarming microscopy experiments in Supplementary Fig. 2, we used a custom enclosed swarming dish. A 22 mm × 22 mm cover glass (with a thickness of 0.13 – 0.17 mm) was sterilized by ethanol and coated with an anti-fog spray (Gamer Advances). 2.5 ml LB medium with 2% agar was solidified in a separate sterile 35-mm Petri dish (Greiner) and allowed to rest at room temperature on the lab bench for 4 hours, until it was moved to 37°C for pre-warming 30 min before the experiment. In the meantime, an 18 mm × 18 mm hole was drilled into the bottom of another 35 mm Petri dish. The edge of this hole was sanded for leveling both interior and exterior surfaces. For sterilization, the dish was soaked in pure ethanol for less than a minute and rinsed thoroughly three times with autoclaved nanopure water. For setting up an experiment, the aforementioned LB agar was carefully removed with a spatula and placed into the Petri dish with the hole. The dish was then closed with a lid and sealed with parafilm. The dish was flipped upside down, and 1 µL of a cell culture was dropped on the agar medium (through the hole). Once the inoculum fully dried out, the hole was closed with a cover glass whose interior surface was coated with anti-fog spray as described above. The cover glass was sealed with Scotch tape. Finally, the dish was moved to a pre-warmed (37°C) microscope.

### WT and Prec competition assay

Liquid cultures with OD_600_ of 1 were prepared as described above for each strain. The resulting cultures were mixed at varying ratios for a final culture volume of 1mL, and 4 µL of the final culture was inoculated onto the center of a pre-warmed swarming agar plate and allowed to dry. Once dry, the plates were moved to an incubator at 37°C with a water reservoir to maintain humidity. After 24 hours of incubation, an inoculum loop was used to collect cells from the edge of each colony. The cells were suspended in 1mL pre-warmed LB and mixed until evenly distributed. The density of the suspension was adjusted to OD_600_ of 0.1, and 100 µL of each culture was inoculated onto LB broth plates with 3% agar to prevent swarming, and sterile glass beads were used to evenly distribute the culture for CFU measurement. Plates were incubated at 37°C for 20 hours before counting the number of colonies with each phenotype.

### Computational modeling

Numerical simulations were performed using custom MATLAB software. We modeled colonies in two dimensions, reduced to one spatial variable by assuming circular symmetry. Solving of the partial differential equations was done using MATLAB’s pdepe solver, which applies a spatial discretization and integrates the resulting ODEs over time to give approximate solutions at specified timepoints [64]. We used a step-function initial condition, and Neumann boundary conditions. Additional details about the models used are provided in the Supplementary Information.

## Acknowledgements

This work was funded by NIH (1U19AI158080, AG, MK) and MP3 Initiative (00097584, ED) We thank Phil Rather for helpful discussions throughout the projects.

## Author Contributions

ED and MK conceived the study. ED and AG designed and carried out the experiments and modeling. MK secured funding and provided resources. ED and MK wrote the manuscript. All authors read and approved the manuscript.

## Data Availability Statements

Source data for figures are provided in the Source Data file.

## Code Availability Statements

Our custom-built software is provided as a Supplementary file.

## Reference

1. Goldschmidt, F., R.R. Regoes, and D.R. Johnson, Successive range expansion promotes diversity and accelerates evolution in spatially structured microbial populations. The ISME Journal, 2017. 11(9): p. 2112–2123.

2. Hill, J.K., H.M. Griffiths, and C.D. Thomas, Climate change and evolutionary adaptations at species’ range margins. Annual review of entomology, 2011. 56: p. 143–59.

3. Tomiolo, S. and D. Ward, Species migrations and range shifts: A synthesis of causes and consequences. Perspectives in Plant Ecology, Evolution and Systematics, 2018.

4. Korolev, K.S., et al., Selective sweeps in growing microbial colonies. Phys Biol, 2012. 9(2): p. 026008.

5. Hastings, A., et al., The spatial spread of invasions: new developments in theory and evidence. Ecology Letters, 2005. 8(1): p. 91–101.

6. Mooney, H.A. and E.E. Cleland, The evolutionary impact of invasive species. Proceedings of the National Academy of Sciences, 2001. 98(10): p. 5446–5451.

7. Sakai, A.K., et al., The Population Biology of Invasive Species. Annual Review of Ecology, Evolution, and Systematics, 2001. 32(Volume 32, 2001): p. 305-332.

8. Lloyd, D.P. and R.J. Allen, Competition for space during bacterial colonization of a surface. Journal of The Royal Society Interface, 2015. 12(110).

9. Korolev, K.S., J.B. Xavier, and J. Gore, Turning ecology and evolution against cancer. Nat Rev Cancer, 2014. 14(5): p. 371–80.

10. Stroud, J.T., et al., Priority effects transcend scales and disciplines in biology. Trends Ecol Evol, 2024. 39(7): p. 677–688.

11. Colautti, R.I. and S.C.H. Barrett, Rapid Adaptation to Climate Facilitates Range Expansion of an Invasive Plant. Science, 2013. 342(6156): p. 364–366.

12. Brown, G.P., C. Kelehear, and R. Shine, The early toad gets the worm: cane toads at an invasion front benefit from higher prey availability. J Anim Ecol, 2013. 82(4): p. 854–62.

13. Duckworth, R.A., Adaptive dispersal strategies and the dynamics of a range expansion. Am Nat, 2008. 172 **Suppl 1**: p. S4–17.

14. Moreau, C., et al., Deep human genealogies reveal a selective advantage to be on an expanding wave front. Science, 2011. 334(6059): p. 1148–50.

15. Vicsek, T., et al., Novel Type of Phase Transition in a System of Self-Driven Particles. Physical Review Letters, 1995. 75(6): p. 1226–1229.

16. Couzin, I.D. and J. Krause, Self-Organization and Collective Behavior in Vertebrates. 2003, Elsevier. p. 1–75.

17. Marchetti, M.C., et al., Hydrodynamics of soft active matter. Reviews of Modern Physics, 2013. 85(3): p. 1143–1189.

18. Kearns, D.B., A field guide to bacterial swarming motility. Nature reviews. Microbiology, 2010. 8(9): p. 634–644.

19. Yan, J., H. Monaco, and J.B. Xavier, The Ultimate Guide to Bacterial Swarming: An Experimental Model to Study the Evolution of Cooperative Behavior. Annual Review of Microbiology, 2019. 73(1): p. 293–312.

20. Partridge, J.D. and R.M. Harshey, More than motility: Salmonella flagella contribute to overriding friction and facilitating colony hydration during swarming. J Bacteriol, 2013. 195(5): p. 919–29.

21. Harshey, R.M., Bees aren’t the only ones: swarming in Gram-negative bacteria. Molecular Microbiology, 1994. 13(3): p. 389–394.

22. Darnton, N.C., et al., Dynamics of bacterial swarming. Biophys J, 2010. 98(10): p. 2082–90.

23. Berg, H.C., Swarming motility: it better be wet. Curr Biol, 2005. 15(15): p. R599–600.

24. Kaiser, D., Bacterial Swarming: A Re-examination of Cell-Movement Patterns. Current Biology, 2007. 17(14): p. R561–R570.

25. Verstraeten, N., et al., Living on a surface: swarming and biofilm formation. Trends Microbiol, 2008. 16(10): p. 496–506.

26. Wu, Y., et al., Self-organization in bacterial swarming: lessons from myxobacteria. Physical Biology, 2011. 8(5): p. 055003.

27. Zhang, H.P., et al., Swarming dynamics in bacterial colonies. EPL (Europhysics Letters), 2009. 87(4).

28. Rørth, P., Collective Cell Migration. Annual Review of Cell and Developmental Biology, 2009. 25(1): p. 407–429.

29. Cheung, K.J. and S. Horne-Badovinac, Collective cell migration modes in development, tissue repair and cancer. Nature Reviews Molecular Cell Biology, 2025. 26(10): p. 741–758.

30. Friedl, P. and D. Gilmour, Collective cell migration in morphogenesis, regeneration and cancer. Nat Rev Mol Cell Biol, 2009. 10(7): p. 445–57.

31. Theveneau, E. and R. Mayor, Collective cell migration of epithelial and mesenchymal cells. Cellular and Molecular Life Sciences, 2013. 70(19): p. 3481–3492.

32. Ouellette, N.T., A physics perspective on collective animal behavior. Phys Biol, 2022. 19(2).

33. Papadopoulou, M., et al., Dynamics of collective motion across time and species. Philosophical Transactions of the Royal Society B, 2023. 378(1874).

34. Toner, J., Y. Tu, and S. Ramaswamy, Hydrodynamics and phases of flocks. Annals of Physics, 2005. 318(1): p. 170–244.

35. May, R.M., Simple mathematical models with very complicated dynamics. Nature, 1976. 261(5560): p. 459–467.

36. Scheffer, M., et al., Catastrophic shifts in ecosystems. Nature, 2001. 413(6856): p. 591–596.

37. Be’er, A. and G. Ariel, A statistical physics view of swarming bacteria. Movement Ecology, 2019. 7(1): p. 9.

38. Hauser, G., Über Fäulnissbacterien und deren Beziehungen zur Septicämie : ein Beitrag zur Morphologie der Speltpilze. 1885, Leipzig: F.C.W. Vogel.

39. Armbruster, C.E. and H.L.T. Mobley, Merging mythology and morphology: the multifaceted lifestyle of Proteus mirabilis. Nature Reviews Microbiology, 2012. 10: p. 743.

40. Simsek, E., et al., Spatial regulation of cell motility and its fitness effect in a surface-attached bacterial community. ISME J, 2022. 16(4): p. 1004–1011.

41. Rauprich, O., et al., Periodic phenomena in Proteus mirabilis swarm colony development. Journal of Bacteriology, 1996. 178(22): p. 6525–6538.

42. Fraser, G.M. and C. Hughes, Swarming motility. Current Opinion in Microbiology, 1999. 2(6): p. 630–635.

43. Alavi, M. and R. Belas, [3] Surface sensing, swarmer cell differentiation, and biofilm development. 2001, Elsevier. p. 29–40.

44. Sturgill, G. and P.N. Rather, Evidence that putrescine acts as an extracellular signal required for swarming in Proteus mirabilis. Mol Microbiol, 2004. 51(2): p. 437–46.

45. Rather, P.N., Swarmer cell differentiation in Proteus mirabilis. Environmental Microbiology, 2005. 7(8): p. 1065–1073.

46. Claret, L. and C. Hughes, Rapid Turnover of FlhD and FlhC, the Flagellar Regulon Transcriptional Activator Proteins, during Proteus Swarming. Journal of Bacteriology, 2000. 182(3): p. 833–836.

47. Clemmer, K.M. and P.N. Rather, Regulation of flhDC expression in Proteus mirabilis. Research in Microbiology, 2007. 158(3): p. 295–302.

48. Furness, R.B., et al., Negative feedback from a Proteus class II flagellum export defect to the flhDC master operon controlling cell division and flagellum assembly. J Bacteriol, 1997. 179(17): p. 5585–8.

49. Murray, J.D., *Mathematical Biology*. Interdisciplinary Applied Mathematics. 2002, New York, NY: Springer New York.

50. Canosa, J., On a Nonlinear Diffusion Equation Describing Population Growth. IBM Journal of Research and Development, 1973. 17(4): p. 307–313.

51. Berry, R.M. and J.P. Armitage, The Bacterial Flagella Motor. 1999, Elsevier. p. 291–337.

52. Lisevich, I., et al., Physics of swimming and its fitness cost determine strategies of bacterial investment in flagellar motility. Nat Commun, 2025. 16(1): p. 1731.

53. Schavemaker, P.E. and M. Lynch, Flagellar energy costs across the tree of life. eLife, 2022. 11.

54. Bonte, D., et al., Costs of dispersal. Biol Rev Camb Philos Soc, 2012. 87(2): p. 290–312.

55. Hatzikirou, H., et al., ’Go or grow’: the key to the emergence of invasion in tumour progression? Math Med Biol, 2012. 29(1): p. 49–65.

56. Smith, J.M. and G.R. Price, The Logic of Animal Conflict. Nature, 1973. 246(5427): p. 15–18.

57. Chase, J.M., Community assembly: when should history matter? Oecologia, 2003. 136(4): p. 489–498.

58. Fukami, T., Historical Contingency in Community Assembly: Integrating Niches, Species Pools, and Priority Effects. Annual Review of Ecology, Evolution, and Systematics, 2015. 46(1): p. 1–23.

59. Couzin, I.D., et al., Collective Memory and Spatial Sorting in Animal Groups. Journal of Theoretical Biology, 2002. 218(1): p. 1–11.

60. Miller, M.B. and B.L. Bassler, Quorum sensing in bacteria. Annu Rev Microbiol, 2001. 55: p. 165–99.

61. Mukherjee, S. and B.L. Bassler, Bacterial quorum sensing in complex and dynamically changing environments. Nat Rev Microbiol, 2019. 17(6): p. 371–382.

62. Worlitzer, V.M., et al., Biophysical aspects underlying the swarm to biofilm transition. Sci Adv, 2022. 8(24): p. eabn8152.

63. Bru, J.-L., A. Siryaporn, and N.M. Høyland-Kroghsbo, Time-lapse Imaging of Bacterial Swarms and the Collective Stress Response. Journal of Visualized Experiments, 2020(159).

64. Skeel, R.D. and M. Berzins, A Method for the Spatial Discretization of Parabolic Equations in One Space Variable. SIAM Journal on Scientific and Statistical Computing, 1990. 11(1): p. 1–32.

