## Supplementary Material for "Early Spatiotemporal Deficits Become Competitive Advantages During Collective Expansion"

**Supplementary Text**

**Section 1: Model for individual diffusivity with logistic growth**

We begin with the classical Fisher–Kolmogorov (FK) equation, which combines a constant rate of diffusion with logistic growth.

$$\frac{\partial u}{\partial t} = D\frac{\partial^2 u}{\partial x^2} + \gamma u(u_{cc} - u), \qquad \text{Eq. S1}$$

where $u(x,t)$ denotes the local population density at position $x$ and time $t$. The parameter $D$ is the diffusion coefficient, representing unbiased individual motility, and $\gamma$ is the intrinsic growth rate. Growth is limited by a carrying-capacity density $u_{cc}$, which sets the maximal sustainable local population density. This formulation assumes that motility is independent of local density and that growth is limited locally by crowding.

By making the substitutions $t' = \gamma t$, $x' = \sqrt{\frac{D}{\gamma}}x$, and $u' = \frac{u}{u_{cc}}$, we nondimensionalize Eq. S1,

$$\frac{\partial u'}{\partial t'} = \frac{\partial^2 u'}{\partial x'^2} + u'(1-u'). \qquad \text{Eq. S2}$$

Prior work has shown that Eq. S2 produces asymptotic travelling wave solutions, where a fixed front profile travels outwards at a constant rate [1, 2]. Here, we are interested in how the colonies with different lag time delays transition to this long-time regime.

We numerically solved Eq. S2. As the initial condition, we prescribed a box-shaped population density profile of height $A$ and width $L$, centered on $x = 0$,

$$u_0(x) = \begin{cases} A & -\frac{L}{2} \le x \le \frac{L}{2} \\ 0 & otherwise \end{cases}. \qquad \text{Eq. S3}$$

The product $V_0 = A\,L$ defines the initial total biomass, which was kept the same across all colonies.

To model delayed initiation, we imposed a waiting period during which diffusion was suppressed ($D = 0$). During this phase, cells proliferate locally, increasing the population density without spatial spreading. After the prescribed waiting time, diffusion was restored, and the population was allowed to expand according to Eq. S2.

The results of the numerical solutions are shown in Fig. 2A. All colonies, regardless of waiting period, converge to a traveling-wave solution with a constant speed, consistent with classical Fisher–Kolmogorov, as expected [1, 2]. Yet, colonies that begin expanding later remain persistently behind by a finite radius offset. Consequently, any initial delay translates directly into a permanent loss in expansion time to a fixed target radius.

**Section 2: Model for collective diffusion with logistic growth**

We extend this framework to collective diffusion, in which effective dispersal is enhanced by local interactions among neighboring individuals. Rather than modeling the details of these interactions explicitly, we capture their net effect phenomenologically by allowing the diffusion coefficient to depend on the local population density, $D = D(u)$, with $D'(u) > 0$.

In this case, Eq. S1 is modified to

$$\frac{\partial u}{\partial t} = \frac{\partial}{\partial x}\left(D(u)\frac{\partial u}{\partial x}\right) + \gamma u(u_{cc} - u). \qquad \text{Eq. S4}$$

Expanding the diffusion operator yields

$$\frac{\partial u}{\partial t} = D(u)\,\frac{\partial^2 u}{\partial x^2} + D'(u)\left(\frac{\partial u}{\partial x}\right)^2 + \gamma u(u_{cc} - u). \qquad \text{Eq. S5}$$

Compared to the constant-diffusion model (Eq. S1), Eq. S5 contains an additional nonlinear term

$$D'(u)\left(\frac{\partial u}{\partial x}\right)^2, \qquad \text{Eq. S6}$$

which arises solely from the density-dependence of diffusion.

Mathematically, the structure of this term implies that spatial gradients in density can directly influence population flux, beyond the effect of random dispersal captured by the linear diffusion term. In particular, when $D'(u) > 0$, regions of higher density contribute to flux along density gradients.

This behavior admits a useful physical interpretation. As a population expands, its density decreases monotonically in the direction of propagation, creating a density gradient. Individuals in denser regions are effectively more motile than those in sparser regions ahead of the front. This asymmetry generates a net flux directed down the density gradient, enhancing forward propagation. Therefore, the nonlinear diffusion term (Eq. S6) can be interpreted as a self-induced, gradient-driven drift.

To clarify this interpretation, it is useful to compare the term with advection in a classical convection–diffusion equation [2], where transport by a velocity field appears as a term proportional to $u\ \partial u/\partial x$. The nonlinear diffusion term (Eq. S6) plays an analogous role: it produces an effective convective bias with a characteristic local "velocity" equal to $D'(u)\ \partial u/\partial x$ [2]. In this sense, collective diffusion can resemble convection.

This mechanism has direct implications for range expansion with delayed initiation. Consider two colonies with the same total biomass: one expands immediately while the other invests a finite waiting period during which diffusion is suppressed while growth proceeds locally. After this period, the colony initiates expansion from a more compact, higher-density profile, which generates steeper density gradients and correspondingly larger values of $|\ \partial u/\ \partial x\ |$. As a result, the nonlinear diffusion term $D'(u)(\partial u/\ \partial x)^2$ is enhanced at early times, producing a stronger, transient outward bias in population flux. Consequently, collective diffusion could reduce—and potentially eliminate—the front-position disadvantage incurred by the lag period.

To test this intuition using numerical simulation, we consider a simple case,

$$D(u) = Du\ , \qquad \text{Eq. S7}$$

where $D$ is a constant. This is the lowest-order density dependence that captures motility enhancement through local interactions and yields a parameter-free nondimensional form.

Substituting $D(u)$ into Eq. S5 gives us

$$\frac{\partial u}{\partial t} = Du\,\frac{\partial^2 u}{\partial x^2} + D\left(\frac{\partial u}{\partial x}\right)^2 + \gamma u(u_{cc} - u). \qquad \text{Eq. S8}$$

By making the substitutions $t' = \gamma t,\ x' = \sqrt{\frac{D}{\gamma}}x$, and $u' = \frac{u}{u_{cc}}$, we nondimensionalize Eq. S8,

$$\frac{\partial u'}{\partial t'} = u'\,\frac{\partial^2 u'}{\partial x'^2} + \left(\frac{\partial u'}{\partial x'}\right)^2 + u'(1 - u'). \qquad \text{Eq. S9}$$

We next performed numerical simulations of this equation using the same box-shaped initial condition and fixed initial total biomass $V_0$ (Eq. S3). As above, delayed initiation was implemented by suppressing diffusion during a prescribed waiting period while allowing growth to proceed locally, after which density-dependent diffusion was restored, and the population was allowed to expand.

The results of simulations are shown in Fig. 2C. Intermediate expansion dynamics somewhat differ from the constant-diffusion FK solutions (Fig. 2A). For example, the no-waiting baseline exhibits slow intermediate expansion, allowing the colonies with delays to reduce the separation. Despite this partial recovery, delayed colonies remain behind. At long times, all trajectories approach a common asymptotic expansion rate but remain separated by a finite radius offset that increases with the duration of the delays. Consequently, even under collective diffusion, delayed initiation results in a persistent expansion-time cost, albeit reduced relative to the individual-diffusion FK case.

This effect is quantified in Fig. 2J, which shows the expansion advantage $S$ as a function of the lag time. Here, $S$ is defined as the difference in arrival time at a fixed target radius relative to the no-waiting baseline (see Fig. 2I); negative values of $S$ therefore indicate a net cost of delayed initiation. In the individual-diffusion FK case, $S$ decreases approximately linearly with lag time, reflecting a persistent arrival-time penalty associated with delayed onset of expansion. Incorporating collective diffusion reduces this cost, yet $S$ remains negative, demonstrating that collective motility mitigates—but does not eliminate—the cost of delayed initiation.

Our additional analysis explains why collective motility does not fully eliminate the cost: under logistic growth, spatially concentrated density profiles intrinsically reduce net biomass production. This can be seen directly by examining the evolution of the total biomass.

Let

$$M(t) = \int u(x,t)\, dx \qquad \text{Eq. S10}$$

denote the total biomass. For logistic growth with a carrying capacity,

$$\frac{dM}{dt} = \int \gamma\, u \left(1 - \frac{u}{u_{cc}}\right) dx = \gamma\, M - \frac{r}{u_{cc}} \int u^2\, dx. \qquad \text{Eq. S11}$$

For a fixed total biomass $M$, a more spatially concentrated distribution has a larger $\int \rho^2\, dx$ and therefore a smaller net growth rate $dM/dt$. Thus, concentrating biomass increases local crowding and suppresses overall biomass accumulation. A colony that delays swarming and remains in lag time will remain more compact than a colony that begins swarming and expands into new territory, and thus the colony with lag time will accumulate biomass more slowly than the swarming colony and develop a deficit in total biomass, as shown in Supplementary Fig. 3.

Once both colonies are swarming and have reached the travelling wave, they will both have the same front profile (with the offset in radius), and any difference in total biomass must be represented behind that front profile in the bulk of the colony, which is effectively a rectangle with a density at the carrying capacity and a radius that is increasing at a constant rate. For the colony with lag time, once it reaches the travelling wave, the total biomass deficit will result in a fixed radius offset behind the colony without lag time, with the size of the radius offset being equal to the total biomass deficit divided by the carrying capacity. This prediction is consistent with our simulation results, which show slower biomass accumulation in colonies with delayed initiation (Supplementary Fig. 3A-B). Therefore, while collective motility mitigates the disadvantage, under logistic growth it does not eliminate it. Consequently, the early head start of the no-delay colony becomes locked in as a finite radius offset.

This observation motivates our consideration of the exponential growth model in the following section, which lacks a carrying capacity and therefore removes the density-dependent growth penalty associated with spatially concentrated population profiles.

**Section 3: Model for individual diffusion with exponential growth**

The equation below describes exponential population growth with constant (density-independent) diffusion:

$$\frac{\partial u}{\partial t} = D \frac{\partial^2 u}{\partial x^2} + \gamma u. \qquad \text{Eq. S12}$$

Nondimensionalizing as discussed above yields

$$\frac{\partial u}{\partial t'} = \frac{\partial^2 u}{\partial x'^2} + u. \qquad \text{Eq. S13}$$

We numerically solved Eq. S13 using the same box-shaped initial condition and fixed initial total biomass $V_0$ (Eq. S3). During this waiting phase, growth proceeds exponentially without saturation, allowing biomass to increase without a density-dependent penalty (Supplementary Fig. 3C). Yet, colonies with delays still fall behind and do not fully catch up to the no-waiting baseline (Fig. 2G).

### Section 4: Model for collective diffusion with exponential growth

To isolate the combined effects of collective motility and unrestricted growth, we replace the logistic growth term in Eq. S12 with exponential growth, yielding

$$\frac{\partial u}{\partial t} = \frac{\partial}{\partial x}\left(D(u)\frac{\partial u}{\partial x}\right) + \gamma u. \qquad \text{Eq. S14}$$

We focus on the minimal form of collective diffusion (Eq. S7). Substituting it into Eq. S14 and nondimensionalizing as described above gives

$$\frac{\partial u}{\partial t'} = \frac{\partial}{\partial x'}\left(u\frac{\partial u}{\partial x'}\right) + u. \qquad \text{Eq. S15}$$

Numerical solutions of Eq. S15 are shown in Fig. 2E and 2F. During the waiting phase, biomass accumulates exponentially without saturation (Supplementary Fig. 3). When diffusion is restored, the resulting high-density, compact profile simultaneously increases diffusion and steepens density gradients, producing self-induced, gradient-driven drift as described in Section 2. As a result, colonies with delays fully recover, converging onto the no-waiting baseline (Fig. 2E).

### Section 5: Reformulating delayed initiation using smaller initial radii

The analyses above show that the cost of delayed initiation arises from the spatial reorganization of biomass that occurs during the waiting period. Suppressing diffusion while allowing growth leads to a more compact, high-density population profile at the moment expansion begins. This compactness alters early expansion dynamics and can enable delayed colonies to recover their initial disadvantage through enhanced collective motility.

We sought to test this prediction experimentally. While the most direct approach would be to artificially impose or remove a waiting period prior to expansion, such delays are often hard-wired by genetic or physiological programs in bacteria and cannot be readily manipulated independently. Instead, we adopt an analogous scenario: colonies that begin expanding simultaneously but differ in their initial spatial extent while having the same total biomass (Supplementary Fig. 4). We justify their equivalence in the main text.

Importantly, as we demonstrate below, this formulation is analytically tractable and provides a transparent geometric interpretation of the delay cost. By isolating spatial compactness as the control parameter, it offers intuitive insight into how early geometric differences translate into persistent expansion advantages or disadvantages.

**Section 5a: Individual diffusion with exponential growth**

We first consider the case of constant diffusion (individual motility) with exponential growth (Eq. S13). For convenience, we drop the prime notation. Eq. S13 can be converted to the standard heat equation [3] by defining a function

$$v(x,t) = e^{-t}u(x,t). \qquad \text{Eq. S16}$$

Substituting into Eq. S13 yields

$$\frac{\partial v}{\partial t} = \frac{\partial^2 v}{\partial x^2}, \qquad \text{Eq. S17}$$

which admits the well-known solution in terms of the heat kernel,

$$v(x,t) = (G_t * u_0)(x) = \int_{-\infty}^{-\infty} \frac{1}{\sqrt{4\pi t}}\, e^{\frac{-(x-y)^2}{4\,t}}\, u_0(y)dy\,, \qquad \text{Eq. S18}$$

where $u_0(x)$ is our initial condition.

Transforming back gives

$$u(x,t) = e^t\, v(x,t) = e^t \int_{-\infty}^{-\infty} \frac{1}{\sqrt{4\pi t}} e^{\frac{-(x-y)^2}{4\,t}} u_0(y)dy\,. \qquad \text{Eq. S19}$$

Incorporating the initial condition (Eq. S3), we have

$$u(x,t) = e^t \int_{-\infty}^{-\infty} \frac{1}{\sqrt{4\pi t}} e^{\frac{-(x-y)^2}{4\,t}} u_0(y)dy \; = e^t \int_{-\frac{L}{2}}^{\frac{L}{2}} \frac{A}{\sqrt{4\pi t}} e^{\frac{-(x-y)^2}{4\,t}} dy = \frac{A}{2} e^t \left( \operatorname{erf}\left( \frac{x+\frac{L}{2}}{\sqrt{4\,t}} \right) - \operatorname{erf}\left( \frac{x-\frac{L}{2}}{\sqrt{4\,t}} \right) \right). \qquad \text{Eq. S20}$$

This solution is consistent with our boundary conditions, $u(\infty,t) = 0$ and $\frac{\partial u(0,t)}{\partial t} = 0$.

Using the definition $V_0 = A\,L$ for the total initial biomass, Eq. S20 can be rewritten as

$$u(x,t) = \frac{V_0}{2\,L} e^t \left( \operatorname{erf}\left( \frac{x+\frac{L}{2}}{\sqrt{4\,t}} \right) - \operatorname{erf}\left( \frac{x-\frac{L}{2}}{\sqrt{4\,t}} \right) \right). \qquad \text{Eq. S21}$$

To analyze the colony expansion, we focus on the dynamics of the far-most front. We choose a small detection threshold density, $\varepsilon$ , defining the front radius, $X(t)$ as

$$u(X(t), t) = \varepsilon. \qquad \text{Eq. S22}$$

Focusing on the rightmost front, we introduce the shifted coordinate for convenience

$$s \coloneqq X(t) - \frac{L}{2} \qquad \text{Eq. S23}$$

We can then make the substitution

$$u(X(t), t) = \frac{V_0}{2\,L} e^t \left( \mathrm{erf}\left( \frac{X(t)+\frac{L}{2}}{\sqrt{4\,t}} \right) - \mathrm{erf}\left( \frac{X(t)-\frac{L}{2}}{\sqrt{4\,t}} \right) \right) = \frac{V_0}{2\,L} e^t \left( \mathrm{erf}\left( \frac{s+L}{\sqrt{4\,t}} \right) - \mathrm{erf}\left( \frac{s}{\sqrt{4\,t}} \right) \right). \qquad \text{Eq. S24}$$

Since we are interested in the rightmost front, we limit ourselves to $s \gg \sqrt{t}$, which allows us to use the Gaussian tail approximation for the difference of the error functions,

$$u(X(t), t) = \frac{V_0}{2\,L} e^t \left( \mathrm{erf}\left( \frac{s+L}{\sqrt{4\,t}} \right) - \mathrm{erf}\left( \frac{s}{\sqrt{4\,t}} \right) \right) \approx \frac{V_0}{L} e^t \left( \frac{1}{s} \sqrt{\frac{t}{\pi}} e^{-\frac{s^2}{4t}} \right). \qquad \text{Eq. S25}$$

Now, we can solve for $s$ using Eq. S22;

$$\frac{V_0}{L} e^t \left( \frac{1}{s} \sqrt{\frac{t}{\pi}} e^{-\frac{s^2}{4t}} \right) = \varepsilon \qquad \text{Eq. S26}$$

$$t - \frac{s^2}{4t} - \log s + \frac{1}{2} \log t + \log\left( \frac{V_0}{L\sqrt{\pi}} \right) = \log \varepsilon \qquad \text{Eq. S27}$$

$$\frac{s^2}{4t} = t + \frac{1}{2} \log t - \log s + \log\left( \frac{V_0}{L\varepsilon\sqrt{\pi}} \right). \qquad \text{Eq. S28}$$

We can then solve for $s$, starting with the ansatz $s = 2t + y(t)$ where $y = o(t)$.

$$\frac{(2t + y)^2}{4t} = t + \frac{1}{2} \log t - \log(2t + y) + \log\left( \frac{V_0}{L\varepsilon\sqrt{\pi}} \right) \qquad \text{Eq. S29}$$

$$t + y + \frac{y^2}{4t} = t + \frac{1}{2}\log t - \log(2t) - \log\left(1 + \frac{y}{2t}\right) + \log\left(\frac{V_0}{L\varepsilon\sqrt{\pi}}\right) \qquad \text{Eq.} S30$$

$$y + \frac{y^2}{4t} = \frac{1}{2}\log t - \log 2 - \log t - \log\left(1 + \frac{y}{2t}\right) + \log\left(\frac{V_0}{L\varepsilon\sqrt{\pi}}\right) \qquad \text{Eq. S31}$$

$$y = -\frac{1}{2}\log t + \log\left(\frac{V_0}{2L\varepsilon\sqrt{\pi}}\right) - \log\left(1 + \frac{y}{2t}\right) - \frac{y^2}{4t}. \qquad \text{Eq. S32}$$

Since $y = o(t)$, $\frac{y^2}{4t} \to 0$ and $\log\left(1 + \frac{y}{2t}\right) = O\left(\frac{y}{t}\right) = o(1)$, giving us

$$y = -\frac{1}{2}\log t + \log\left(\frac{V_0}{2L\varepsilon\sqrt{\pi}}\right) + \mathrm{o}(1)\ . \qquad \text{Eq. S33}$$

We now have

$$s = 2t + y(t) = 2t - \frac{1}{2}\log t + \log\left(\frac{V_0}{2L\varepsilon\sqrt{\pi}}\right) + \mathrm{o}(1)\,, \qquad \text{Eq. S34}$$

which we can substitute into Eq S23 to get the front position

$$X(t) = \frac{L}{2} + 2t - \frac{1}{2}\log t + \log\left(\frac{V_0}{2\ L\ \varepsilon\sqrt{\pi}}\right) + o(1) \qquad \text{Eq. S35}$$

for large $t$.

We can then compare the front location for two different initial radii where $L_1 > L_2$,

$$X_1(t) = \frac{L_1}{2} + 2t - \frac{1}{2}\log t + \log\left(\frac{V_0}{2\ L_1\ \varepsilon\sqrt{\pi}}\right) + o(1) \qquad \text{Eq. S36}$$

$$X_2(t) = \frac{L_2}{2} + 2t - \frac{1}{2}\log t + \log\left(\frac{V_0}{2\ L_2\ \varepsilon\sqrt{\pi}}\right) + o(1). \qquad \text{Eq. S37}$$

Their difference is

$$X_1(t) - X_2(t) = \left(\frac{L_1}{2} + 2t - \frac{1}{2}\log t + \log\left(\frac{V_0}{2\ L_1\ \varepsilon\sqrt{\pi}}\right) + o(1)\right) - \left(\frac{L_2}{2} + 2t - \frac{1}{2}\log t + \log\left(\frac{V_0}{2\ L_2\ \varepsilon\sqrt{\pi}}\right) + o(1)\right) \quad \text{Eq. S38}$$

$$X_1(t) - X_2(t) = \frac{L_1 - L_2}{2} + \log\left(\frac{L_2}{L_1}\right) + o(1). \qquad \text{Eq. S39}$$

Therefore, at late times, the edges of the two colonies will approach a fixed separation that depends on the initial radius of the colony. In particular, a colony that begins from a smaller initial radius never catches up to one that begins from a larger radius, even though the total biomass is identical. This result demonstrates that under exponential growth with constant diffusion, initial geometric differences lead to persistent expansion advantages.

**Section 5b: Collective diffusion with exponential growth**

Starting from Eq. S15, we introduce the transformations

$$v(x,t) = e^{-t}u(x,t)\,, \tau = e^{t} - 1, \qquad \text{Eq. S40}$$

$v$ satisfies the Porous medium equation [3]

$$\frac{\partial v}{\partial \tau} = \frac{\partial}{\partial x}\left(v\,\frac{\partial v}{\partial x}\right). \qquad \text{Eq. S41}$$

While we can't solve this analytically for all time, we can rely on the self-similar Barenblatt solution to the Porous medium equation [4], giving us

$$v(x,\tau) = \tau^{-\frac{1}{3}}\left(C - \frac{x^2}{6\,\tau^{\frac{2}{3}}}\right). \qquad \text{Eq. S42}$$

The constant $C$ is determined by conservation of total mass. Using the initial condition of a box of height $A$ and width $L$, mass conservation gives

$$AL = \int_0^{R(\tau)} \tau^{-\frac{1}{3}}\left(C - \frac{x^2}{6\,\tau^{\frac{2}{3}}}\right)dx = \frac{2\sqrt{6}}{3}C^{\frac{3}{2}}, \qquad \text{Eq. S43}$$

$$C = \left(\frac{3AL}{2\sqrt{6}}\right)^{\frac{2}{3}}. \qquad \text{Eq. S44}$$

Finally, we transform back to $u$ to get the asymptotic solution

$$u(x,t) \sim e^{t}(e^{t}-1)^{-\frac{1}{3}}\left(C - \frac{x^2}{6\,(e^{t}-1)^{\frac{2}{3}}}\right). \qquad \text{Eq. S45}$$

At large $t$, we can further simplify to

$$u(x,t) \sim e^{\frac{2t}{3}}\left(C - \frac{x^2}{6\,e^{\frac{2t}{3}}}\right) = e^{\frac{2t}{3}}\left(\left(\frac{3AL}{2\sqrt{6}}\right)^{\frac{2}{3}} - \frac{x^2}{6\,e^{\frac{2t}{3}}}\right). \qquad \text{Eq. S46}$$

Using the total initial biomass, $V_0 = A * L$, we can write

$$u(x,t) \sim e^{\frac{2t}{3}}\left(\left(\frac{3V_0}{2\sqrt{6}}\right)^{\frac{2}{3}} - \frac{x^2}{6\,e^{\frac{2t}{3}}}\right). \qquad \text{Eq. S47}$$

Crucially, the asymptotic solution depends only on the total initial biomass $V_0$ and not on the initial spatial extent of the colony. Consequently, colonies with the same total biomass but different initial radii converge onto the same asymptotic profile at long times. This means that colonies that begin from smaller initial radii fully catch up to a larger one.

### Section 6: Optimizing time to occupation to predict the threshold-like initiation of collective motility

In the main text, we showed that under collective motility, delaying expansion can provide a competitive advantage. That analysis, however, was based on the asymptotic travelling-wave regime and is therefore most relevant to expansion in effectively unbounded space. In finite habitats, the relevant quantity is instead the total time required for a population to reach a boundary.

We therefore define the total time to occupation as

$$T_{\text{total}}(u) = T_L(u) + T_V(u), \qquad \text{Eq. S48}$$

where $T_L(u)$ is the lag time required for the population to grow from the initial inoculum density $u_0$ to the initiation density $u$, and $T_V(u)$ is the subsequent travel time required to reach the boundary once expansion begins.

#### Section 6a: Lag time

Assuming exponential growth during the lag phase,

$$u(t) = u_0 e^{\lambda_{\text{lag}} t}, \qquad \text{Eq. S49}$$

the lag time required to reach density $u$ is

$$T_L(u) = \frac{1}{\lambda_{\text{lag}}} \ln\left(\frac{u}{u_0}\right). \qquad \text{Eq. S50}$$

Thus, $T_L(u)$ is a monotonically increasing function of $u$: reaching a higher initiation density requires waiting longer. The slope of this increase is set by $\lambda_{\text{lag}}$, so faster growth during lag reduces the temporal cost of waiting.

#### Section 6b: Travel time

The benefit of waiting is that a larger initiation density enhances subsequent expansion. If the boundary is located at distance $L$, we write the travel time as

$$T_V(u) = \frac{L}{v(u)}, \qquad \text{Eq. S51}$$

where $v(u)$ is the effective expansion velocity after initiation. Under collective motility, $v(u)$

increases with $u$, because higher density enhances transport. Therefore, $T_V(u)$ is a decreasing function of $u$.

The precise form of $v(u)$ depends on the precise transient dynamics prior to the asymptotic regime. For the present argument, only the monotonic dependence matters: increasing the initiation density reduces the subsequent travel time.

The total time to occupation is therefore shaped by two opposing effects. Increasing $u$

1. increases $T_L(u)$, because the population must wait longer before expansion begins, and
2. decreases $T_V(u)$, because expansion proceeds faster from a denser initial state.

**Section 6c: Existence of a minimum**

Because the lag term grows without bound,

$$T_L(u) = \frac{1}{r}\ln\left(\frac{u}{u_0}\right) \to \infty \;\text{ as } u \to \infty, \qquad \text{Eq. S52}$$

the total occupation time also diverges at large $u$,

$$T_{\text{total}}(u) \to \infty. \qquad \text{Eq. S53}$$

Thus, $T_{\text{total}}(u)$ must attain a minimum for $u \geq u_0$.

This minimum may be either the trivial boundary optimum $u^* = u_0$, corresponding to immediate initiation without delay, or a nontrivial interior optimum $u^* > u_0$.

**Section 6d: Condition for a nontrivial optimal initiation density**

An interior optimum $u^*$ is determined by minimizing $T_{\text{total}}(u)$:

$$\frac{dT_{\text{total}}}{du} = \frac{1}{\lambda_{\text{lag}} u} + \frac{dT_V}{du}. \qquad \text{Eq. S54}$$

At an interior minimum,

$$\frac{dT_{\text{total}}}{du}\Big|_{u=u^*} = 0 \Leftrightarrow \frac{dT_V(u^*)}{du} = -\frac{1}{\lambda_{\text{lag}} u^*}. \qquad \text{Eq. S55}$$

For such a point to exist, the derivative $dT_{\text{total}}/du$ must become negative over some interval of densities before eventually turning positive again at large $u$. This requires that the decrease in travel time with increasing density be strong enough to outweigh the waiting cost:

$$\frac{dT_V}{du} < -\frac{1}{\lambda_{\text{lag}} u} \text{ for some } u > u_0. \qquad \text{Eq. S56}$$

**Section 6e: Role of growth rate and collective motility**

Equation S56 clarifies how biological parameters influence the existence of a nontrivial optimal initiation density. First, a larger lag-phase growth rate $\lambda_{\text{lag}}$ makes the right-hand side,

$$-\frac{1}{\lambda_{\text{lag}} u}, \quad \text{Eq. S57}$$

closer to zero. This weakens the temporal penalty of waiting and makes the condition for an interior optimum easier to satisfy.

Second, collective motility causes travel time to decrease more steeply with initiation density, making $dT_V/du$ more negative. This strengthens the benefit of waiting.

In summary, fast growth during the lag phase and strong density-dependent enhancement of expansion both favor the emergence of a nontrivial optimal initiation density. At this optimal density, the total time required to occupy the habitat is minimized.

**Section 6f. Inferring the initiation density from lag–inoculum measurements**

To test whether swarming is initiated upon reaching a critical density, we inferred the initiation density $u_c$ from the measured relationship between lag time and initial inoculum density (Fig. 5). The density during lag increases

$$u(t) = u_0 e^{\lambda_{\text{lag}} t}. \quad \text{Eq. S58}$$

If expansion begins when the population reaches the initiation density $u_c$, then the lag time $T_L$ satisfies

$$u_c = u_0 e^{\lambda_{\text{lag}} T_L}. \quad \text{Eq. S59}$$

Solving for $T_L$ gives

$$T_L = \frac{1}{\lambda_{\text{lag}}} \ln\left(\frac{u_c}{u_0}\right). \quad \text{Eq. S60}$$

Thus, if $u_c$ is constant, the lag time should vary linearly with $-\ln\ u_0$, with slope $-1/\lambda_{\text{lag}}$.

For each species, we fit the measured lag times as a function of initial inoculum density using the log-linear form above. From the fitted slope, we estimated the effective growth rate during lag as

$$\lambda_{\text{lag}} = -\frac{1}{\text{slope}}. \quad \text{Eq. S61}$$

Using the fitted growth rate $\lambda_{\text{lag}}$ and the measured lag time $T_L$ for each inoculum condition, we inferred the initiation density as

$$u_c = u_0 e^{\lambda_{\text{lag}} T_L}. \qquad \text{Eq. S62}$$

The results are plotted as Supplementary Fig. 7.

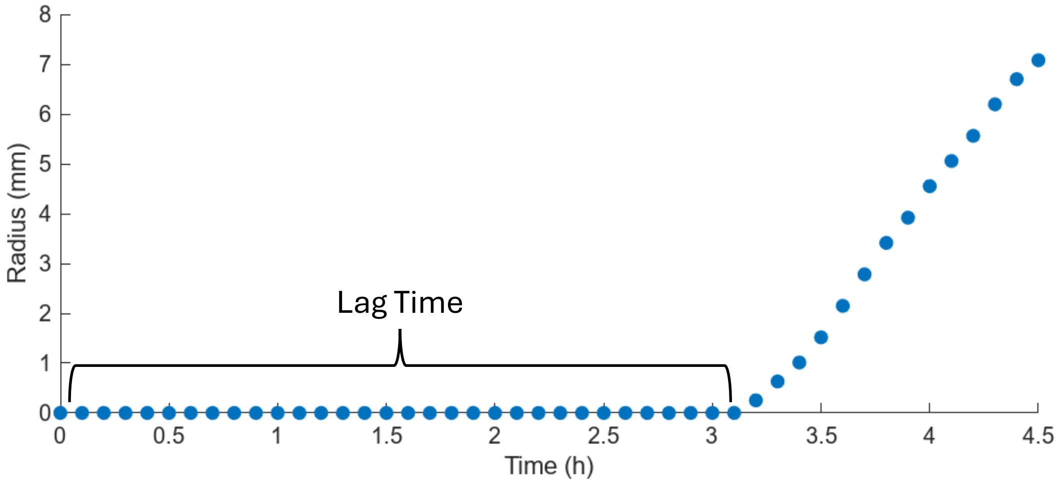


**Supplementary Figure 1. Onset of swarming is delayed in *P. mirabilis*.**

We inoculated *P. mirabilis* on a hard surface (time zero) and recorded the time-lapse images using a scanner. After inoculation, it exhibits a period of lag time (bracket). Following the lag time, the colony expansion begins, which is characterized by coordinated radial movement of the colony boundary.

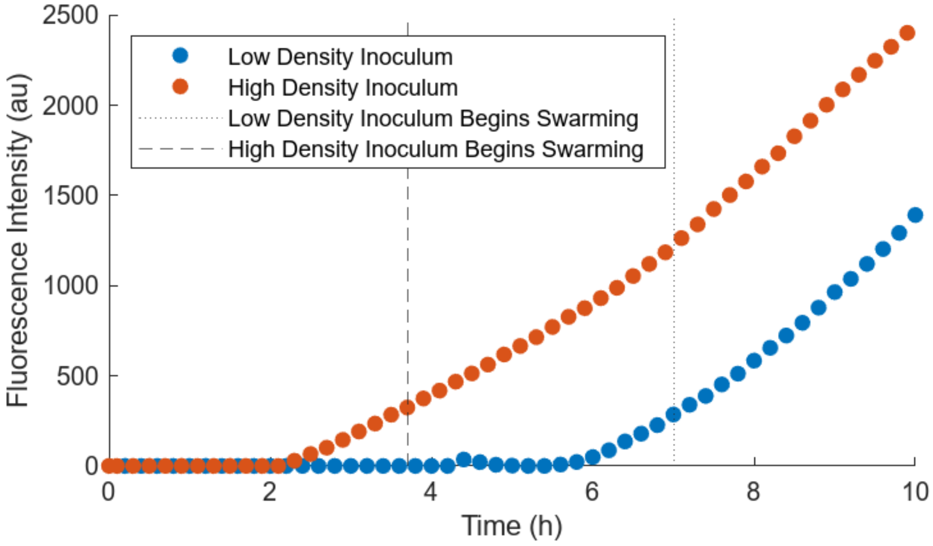


**Supplementary Figure 2. *flhDC* upregulation**

To probe the role of active regulation in the density dependent lag times observed in Fig. 1, we monitored expression of *flhDC*, a master regulator whose upregulation triggers swarming [5]. We used a strain with a transcriptional fusion of the *gfpmut3b* gene to the native *flhDC* operon [6]. We compared a high-density inoculum ($OD_{600}$=10) to a low-density inoculum ($OD_{600}$=0.001), both grown on swarming agar plates and observed using both phase contrast (to monitor radial expansion) and fluorescence (to monitor *flhDC* expression) microscopy. For each colony, we measured the average fluorescence intensity in a region of approximately 200μm × 200μm located near the edge of the original inoculum. The high-density inoculum exhibited early induction of *flhDC*, followed by swarming initiation (vertical dashed line). In contrast, the low-density inoculum displayed delayed *flhDC* activation and correspondingly delayed swarming onset (vertical dotted line). These results indicate that swarming initiation is controlled by density-dependent gene regulation.

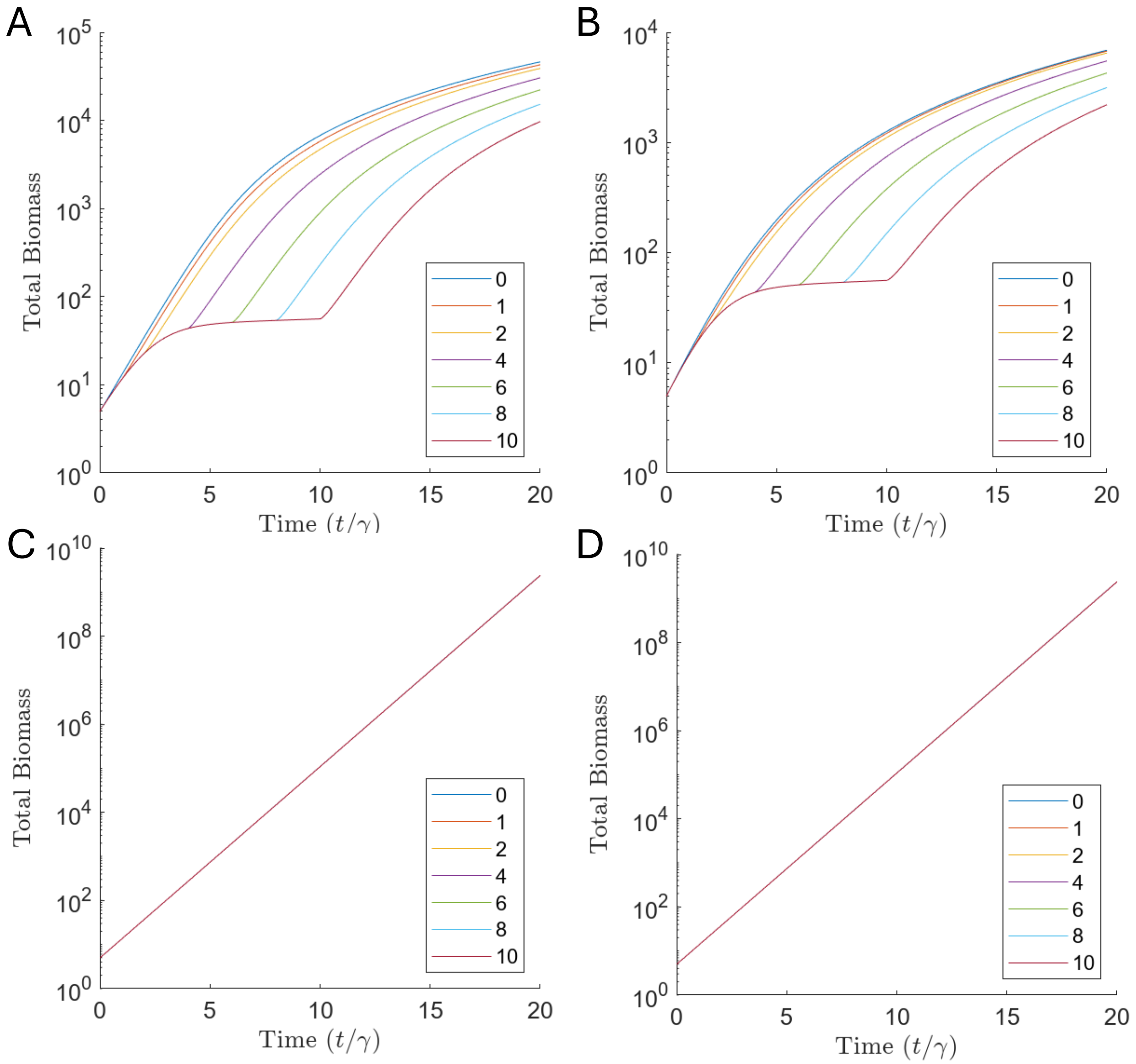


**Supplementary Figure 3. Under logistic growth, increased lag time impedes total biomass accumulation.**

The total colony biomass was calculated for modeled colonies with varied lag times under A. individual motility, logistic growth, B. collective motility, logistic growth, C. individual motility, exponential growth, and D. cooperative motility, exponential growth models.

For logistic growth (A, B), in both the individual motility and collective motility cases, colonies remain more compact during their lag time, leading to lower net growth rates and slower accumulation of biomass. Even after swarming begins, the delayed colonies cannot make up the difference in biomass, ending up with a fixed biomass deficit once all colonies have reached the travelling wave. For exponential growth (C, D), net colony growth rates are independent of colony shape, so no difference in total biomass develops.

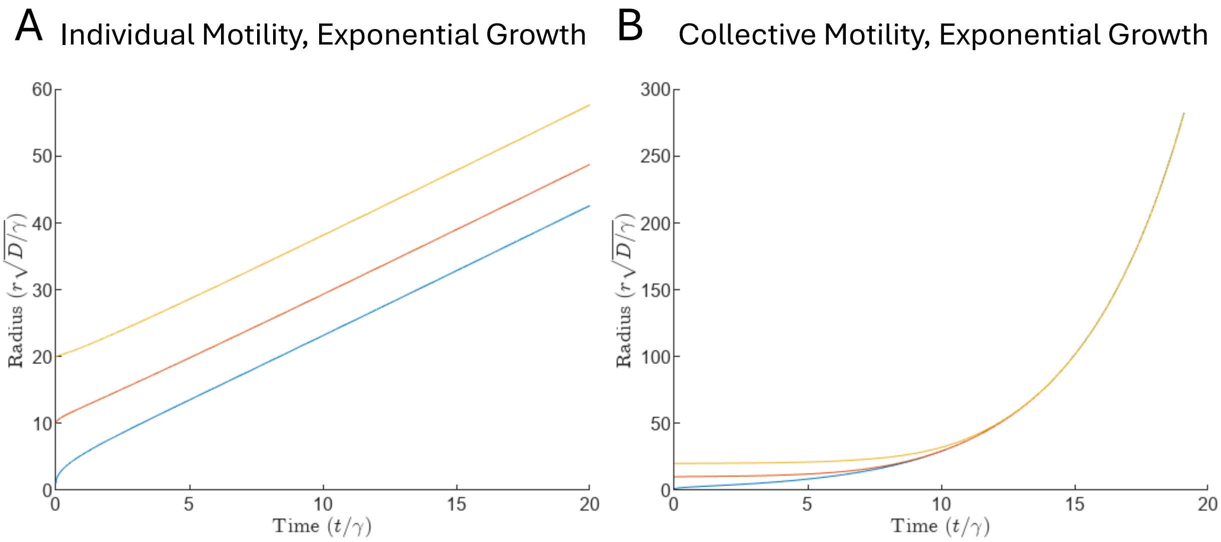


**Supplementary Figure 4. Collective motility mitigates sensitivity to initial conditions.**

Colonies with the same initial biomass and different initial radii were modeled under A. individual motility, exponential growth and B. collective motility, exponential growth models. As with the colonies modeled with identical initial conditions but varied lag times (Fig. 2E,G), in the individual motility, exponential growth case colonies with smaller initial radii never catch up to colonies with larger initial radii, but in the collective motility, exponential growth case, all colonies reach the same radius by the time they reach the travelling wave.

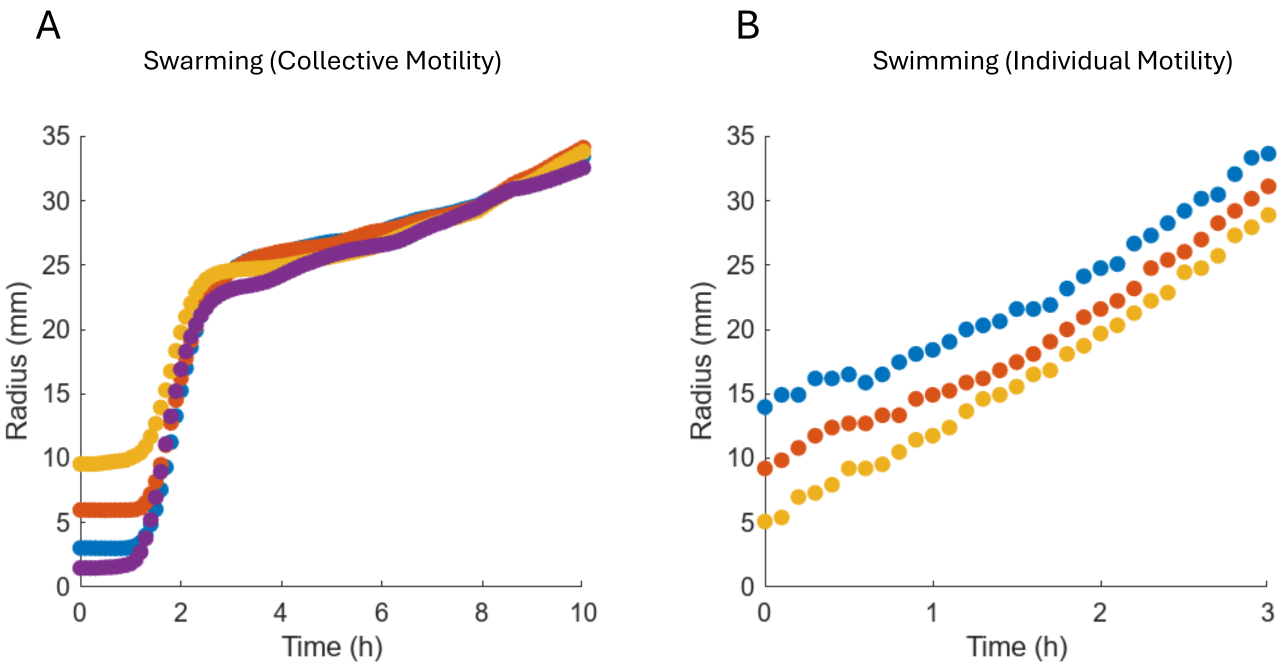


**Supplemental Figure 5. Repeated experimental data shows collective motility mitigates sensitivity to initial conditions.**

We performed a biological repeat of the swimming and swarming experiments in Fig. 3. Swimming and swarming agar plates were inoculated with the same total number of cells, but different initial radii. In the swarming (collective motility) case (A), colonies that start with a smaller radius catch up to colonies that start with a larger initial radius, while in the swimming (individual motility) case (B), colonies that start with a larger radius maintain that advantage as they expand.

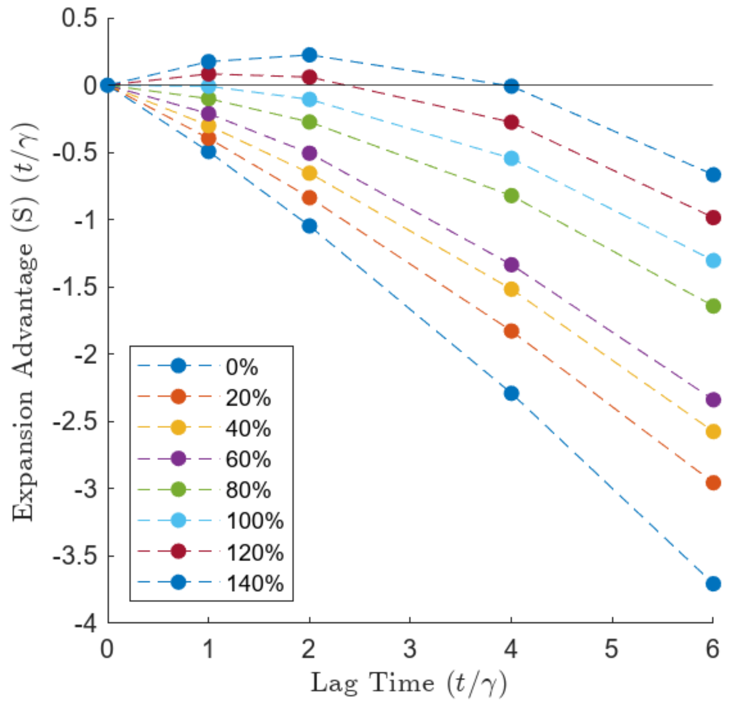


**Supplementary Figure 6. With a growth-motility tradeoff, a positive expansion advantage is sometimes possible with individual motility.**

In the main text, we showed the expansion advantage, *S*, for collective motility (and exponential growth); see Fig, 2I,J. Here, we repeated this analysis for individual motility. We found that, while it is possible to achieve a positive growth advantage (S), it requires a significant boost to lag time growth rate, a ~120% increase relative to the growth rate during movement. This required degree of growth rate boost is much higher than what we observed experimentally in *P. mirabilis* (Fig. 4A).

A

| Strain | Surface Growth Rate (h$^{-1}$) |
|---|---|
| P. mirabilis | 2.6 |
| B. subtilis | 2.9 |
| E. coli | 1.4 |
| P. aeruginosa | 1.7 |
| Evolved P. mirabilis | 3.0 |

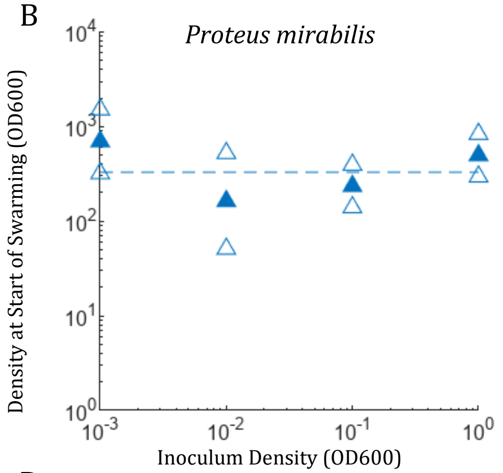


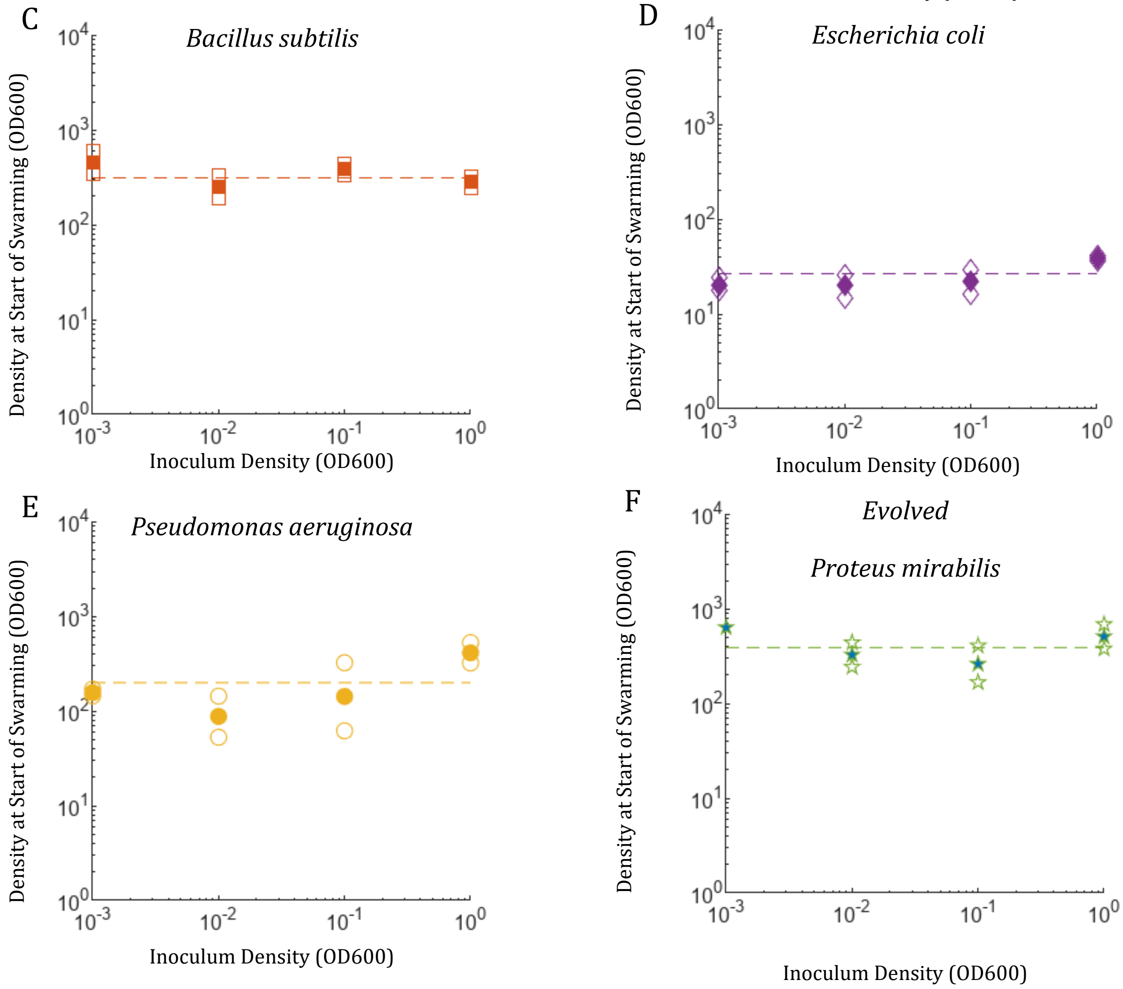

G

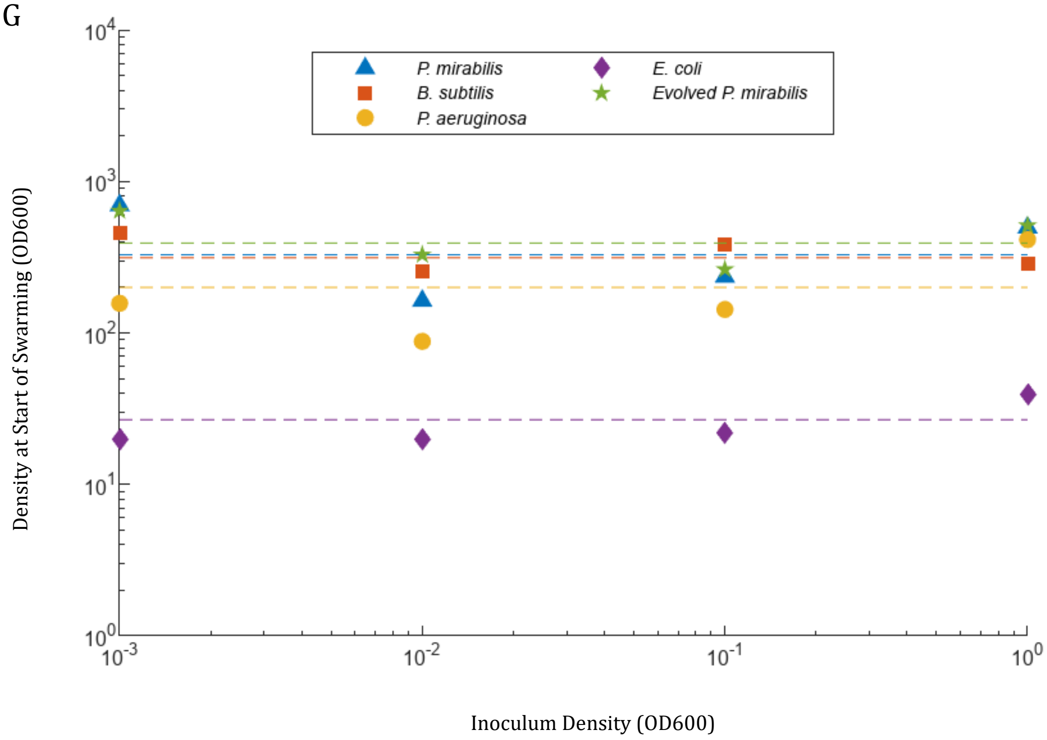


**Supplementary Figure 7. Density at the start of swarming remains relatively constant across inoculum density.**

Using the lag-inoculum data in Fig. 5, we infer (A) the surface growth rate, and (B-G) the inoculum density of each colony at the onset of swarming ($u_c$) (see Supplementary Text, Section 6f). Open symbols indicate individual replicates, closed symbols indicate the mean. For each strain, $u_c$ remains relatively constant across initial inoculum density.

**Supplementary Video 1 | Delayed initiation under individual motility with logistic growth.**

Simulation of range expansion with density-independent motility and logistic growth for colonies with identical initial biomass, with and without an imposed lag phase. During the lag phase, the delayed colony does not expand. After motility is restored, colonies follow nearly identical expansion dynamics shifted in time, resulting in a permanent spatial offset. Delayed colonies therefore remain behind at long times (Fig. 2A,B).

**Supplementary Video 2 | Delayed initiation under collective motility with logistic growth.**

Simulation of range expansion with density-dependent collective motility and logistic growth for colonies with identical initial biomass, with and without an imposed lag phase. Delayed colonies undergo a transient acceleration after the onset of expansion, leading to partial catch-up. However, this speed boost is insufficient to erase the initial disadvantage, and a residual spatial offset remains at long times (Fig. 2C,D).

**Supplementary Video 3A | Short lag under collective motility with exponential growth.**

Simulation of range expansion with density-dependent collective motility and exponential growth for colonies with identical initial biomass. A lag generates a transient acceleration after the onset of expansion, allowing the delayed colony to catch up fully to the no-delay reference. This illustrates that under collective motility in the low-density regime, delayed initiation carries no long-term cost (Fig. 2E,F).

**Supplementary Video 3B | Long lag under collective motility with exponential growth.**

Same as Supplementary Video 3A, but with a longer imposed lag. Despite the larger initial disadvantage, the delayed colony still fully catches up through a stronger transient acceleration after expansion begins. Even long delays therefore remain cost-free under collective motility with exponential growth.

**Supplementary Video 4 | Delayed initiation under individual motility with exponential growth.**

Simulation of range expansion with density-independent motility and exponential growth for colonies with identical initial biomass, with and without an imposed lag phase. Although growth remains exponential, delayed colonies still fail to catch up and retain a permanent spatial offset, showing that exponential growth alone is insufficient to eliminate the cost of delay (Fig. 2G,H).

**Supplementary Table 1. Strains used in this study.**

| Strain | Description/Genotype | Derived from | Comment | Reference |
|---|---|---|---|---|
| ***P. mirabilis* strains** | | | | |
| AMK3 | -325 *flhDC*::KanR | AMK50 | A mini-Tn*5lacZ* insertion 325 bp upstream of the *flhDC* transcriptional start site | [5, 7] |
| AMK5 | Δ*motB*::genR | AMK50 | | Originally from the laboratory of Philip N. Rather |
| AMK50 | $Tc^R$ | - | PM7002 ATCC wild-type strain | Originally from the laboratory of Philip N. Rather |
| AMK79 | $Tc^R$ *flhDC*-*gfpmut3b*-$Amp^R$ | AMK50 | Transcriptional fusion of the *gfpmut3b* gene to the native *flhDC* operon | [6] |
| ***B. subtilis* strains** | | | | |
| AMK42 | | | NCIB 3610 wild-type strain | Originally from the laboratory of Daniel Kearns |
| ***E. coli* strains** | | | | |
| EMK7 | | | AW405 Chemotactic wild-type strain | |
| ***P. aeruginosa* strains** | | | | |
| AMK9 | | | PAO1 wild-type strain | Originally from the laboratory of Joanna Goldberg |

**Supplementary References**


1. Canosa, J., *On a Nonlinear Diffusion Equation Describing Population Growth.* IBM Journal of Research and Development, 1973. **17**(4): p. 307-313.

2. Murray, J.D., *Mathematical Biology*. Interdisciplinary Applied Mathematics. 2002, New York, NY: Springer New York.

3. Evans, L.C., *Partial Differential Equations*. Second Edition ed. Graduate Studies in Mathematics. Vol. 19. 2010: American Mathematical Society.

4. Barenblatt, G.I., *On self-similar motions of compressible fluid in a porous medium.* Prikladnaya Matematika i Mekhanika (Applied Mathematics and Mechanics (PMM)), 1952. **16**(6): p. 679-698.

5. Clemmer, K.M. and P.N. Rather, *Regulation of flhDC expression in Proteus mirabilis.* Research in Microbiology, 2007. **158**(3): p. 295-302.

6. Simsek, E., et al., *Spatial regulation of cell motility and its fitness effect in a surface-attached bacterial community.* ISME J, 2022. **16**(4): p. 1004-1011.

7. Sturgill, G. and P.N. Rather, *Evidence that putrescine acts as an extracellular signal required for swarming in Proteus mirabilis.* Mol Microbiol, 2004. **51**(2): p. 437-46.
